# Integrative Analysis of Epilepsy-Associated Genes Reveals Expression-Phenotype Correlations

**DOI:** 10.1101/2023.06.09.544142

**Authors:** Wanhao Chi, Evangelos Kiskinis

## Abstract

Epilepsy is a highly prevalent neurological disorder characterized by recurrent seizures. Patients exhibit broad genetic, molecular, and clinical diversity involving mild to severe comorbidities. The factors that contribute to this phenotypic diversity remain unclear. We used publicly available datasets to systematically interrogate the expression pattern of 247 epilepsy-associated genes across human tissues, developmental stages, and central nervous system (CNS) cellular subtypes. We grouped genes based on their curated phenotypes into 3 broad classes: core epilepsy genes (CEG), where seizures are the core syndrome, developmental and epileptic encephalopathy genes (DEEG) that are associated with developmental delay, and seizure-related genes (SRG), which are characterized by developmental delay and gross brain malformations. We find that DEEGs are highly expressed within the CNS, while SRGs are most abundant in non-CNS tissues. DEEGs and CEGs exhibit highly dynamic expression in various brain regions across development, spiking during the prenatal to infancy transition. Lastly, the abundance of CEGs and SRGs is comparable within cellular subtypes in the brain, while the average expression level of DEEGs is significantly higher in GABAergic neurons and non-neuronal cells. Our analysis provides an overview of the expression pattern of epilepsy-associated genes with spatiotemporal resolution and establishes a broad expression-phenotype correlation in epilepsy.

## Introduction

Epilepsy constitutes one of the most common neurological disorders estimated to affect 70 million people worldwide^1^. It is primarily characterized by recurrent and unprovoked seizures in the brain^2^. Epilepsy patients often exhibit developmental impairments and other comorbidities^3^. The pathophysiological mechanisms underlying the phenotypic diversity of epilepsy patients remain unclear.

Genetic predisposition plays an important role in epilepsy^4^, and several hundred epilepsy-associated genes have been identified^5,6^. These genes have diverse molecular functions and are implicated in different disease subtypes. One intriguing question is whether the diverse clinical manifestation seen in patients correlates with the spatiotemporal expression of associated genes. This question has not been systematically addressed, partially due to the lack of extensive human gene expression datasets. In recent years, several large and publicly-accessible gene expression data have been assembled including complete human tissue expression data from the Genotype-Tissue Expression (GTEx) project^7^, and brain developmental and cell type-specific expression data from the Allen Brain Institute^8,9^. The availability of these extensive and high-quality datasets makes it possible to investigate gene expression-phenotype correlations.

Here we examined the expression pattern of epilepsy-associated genes at the tissue, developmental stage, and single-cell levels. We used a list of 247 epilepsy-associated genes from the Online Mendelian Inheritance in Man (OMIM)^10^ catalog that records gene-disease relationships, and categorized these genes into three broad groups based on their associated phenotypes: a) <u>c</u>ore <u>e</u>pilepsy <u>g</u>enes (CEG), b) <u>d</u>evelopmental and <u>e</u>pileptic <u>e</u>ncephalopathy <u>g</u>enes (DEEG), and c) <u>s</u>eizure-related <u>g</u>enes (SRG). To examine their expression, we obtained bulk RNA sequencing (RNA-seq) and single nucleus RNA-seq datasets from GTEx^7^ and the Allen Brain Atlas^8,9^ and performed hierarchical clustering analyses. We found that the CEGs and DEEGs exhibit differential expression between central nervous system (CNS) and non-CNS tissues, that DEEGs are expressed 2-to-3-fold higher than CEGs or SRGs in various brain regions across development, and that expression of individual epilepsy-associated genes is largely comparable between cortical GABAergic and glutamatergic neurons. The high expression level of SRGs in non-CNS tissues and of DEEGs during development may contribute to non-CNS comorbidities and neurodevelopmental impairments that are associated with these two classes of genes respectively. Our analysis provides an overview of the expression pattern of epilepsy-associated genes with spatiotemporal resolution and establishes a broad expression-phenotype correlation in epilepsy.

## Methods

### Categorization of epilepsy-associated genes based on phenotypes

The genotype-phenotype data were obtained from OMIM (accessed in November, 2022)^10^. A total of 8533 items were curated in OMIM, including disorders with unknown genetic defects (n = 33), disorders with genes but no mutation identified (n = 1051), disorders with genes and mutations identified (n = 7285), and disorders with multiple genes involved through deletion or duplication (n = 164). We queried related items from the group of disorders with genes and mutations identified using the following terms: [E/e]pilepsy, [E/e]pilepsies, [E/e]pileptic, [S/s]eizure, and [S/s]eizures, which led to 282 items and 247 unique genes.

### Functional classification of epilepsy-associated genes

Functional classification of epilepsy-associated genes was performed using the PANTHER classification system^11^. Based on assigned PANTHER Protein Class, we grouped genes with known functions to one of the following seven categories: enzyme, ion channel, transporter/receptor, signaling, transcription/translation, cytoskeleton, and extracellular matrix. Genes with unknown functions were grouped into the category of ‘unknown‘.

### Expression of epilepsy-associated genes in human tissues

Tissue-specific transcriptomes were obtained from the Genotype-Tissue Expression (GTEx) project (Open access data, V8 dbGaP Accession phs000424.v8.p2)^7^. This dataset consists of bulk RNA sequencing data from 663 donors across 52 tissues as well as cultured fibroblast cells and transformed lymphocytes. For the clustering analyses we considered the mean expression for each gene across 52 tissues. The final data included 247 epilepsy-associated genes.

### Expression of epilepsy-associated genes in the brain across development

Developmental transcriptomes in the brain were obtained from BrainSpan (RNA-Seq Gencode v10 summarized to genes)^9^. This dataset consists of bulk RNA sequencing data in 26 brain sub-regions in 11 developmental stages, from embryonic stages to adulthood, with the sample size of each region ranging from 229 to 8244 in total. We selected epilepsy-associated genes in this dataset and grouped every single stage into one of five periods: prenatal (4-37 post conception weeks), infancy (birth-18 months), childhood (19 months-11 years), adolescence (12-19 years), and adulthood (20-60+ years), based on the criteria used by BrainSpan^9^. We analyzed 229 epilepsy-associated genes that had expression data across all five developmental periods in 16 brain sub-regions. Brain regions with no data for all periods were excluded.

### Expression of epilepsy-associated genes in brain cell types

Brain cell-type specific transcriptomes were obtained from BrainMap^8^. This dataset consists of single-nucleus RNA sequencing data in 49,474 nuclei from six areas of the human cortex (primary motor cortex, primary sensory cortex, primary auditory cortex, primary visual cortex, middle temporal gyrus, and anterior cingulate gyrus). We selected individual cells that are molecularly defined (i.e., glutamatergic, GABAergic, or non-neuronal cells) and exhibit expression of epilepsy-associated genes. The final analyses include 236 epilepsy-associated genes in 45 clusters from 10,620 single nuclei.

### Data analysis and plotting

Analyses were performed in R (www.R-project.org). The Circos plot and heatmaps were generated using the Circlize^12^ and Complex Heatmap package^13^. The R packages and codes for data analysis are available at Github.

## Results

### Selection and categorization of epilepsy-associated genes into three distinct groups based on clinical phenotype

We obtained 247 epilepsy-associated genes from OMIM, an Online Catalog of Human Genes and Genetic Disorders that curates both genes and mutations with related clinical phenotypes in patients^10^ (Supplemental Table 1). Based on the curated phenotypes, we categorized genes into three groups: a) core epilepsy genes (CEG), b) developmental and epileptic encephalopathy genes (DEEG), and c) seizure-related genes (SRG) (Fig. 1, Supplemental Table 1). CEGs are genes identified in epilepsy patients with seizures as the core syndrome. DEEGs are genes identified in patients with developmental and epileptic encephalopathy (DEE). According to the definition from the International League Against Epilepsy (ILAE), DEE patients have gradually increased cognitive and behavioral impairments caused by epileptic activity^3^. Notably, DEE patients do not typically exhibit gross brain malformation, which contrasts with SRG-associated diseases, where patients often show brain malformation or other developmental defects^3^. While most genes are unique within each group, we found thirteen genes shared by CEG and DEEG: *GABRA1, GABRB3, SCN3A, SLC12A5, TBC1D24, SCN1A, GABRG2, SCN1B, HCN1, SCN2A, SCN8A, KCNQ2*, and *KCNT1*. We grouped these genes within CEGs for clarity. Additionally, *KCNMA1* is associated with both CEG and SRG and we grouped it within CEGs, while *ATP6V0A1, GRIN1*, and *GRIN2B* are associated with both DEEG and SRG and we included them within DEEGs.

**Figure 1.**
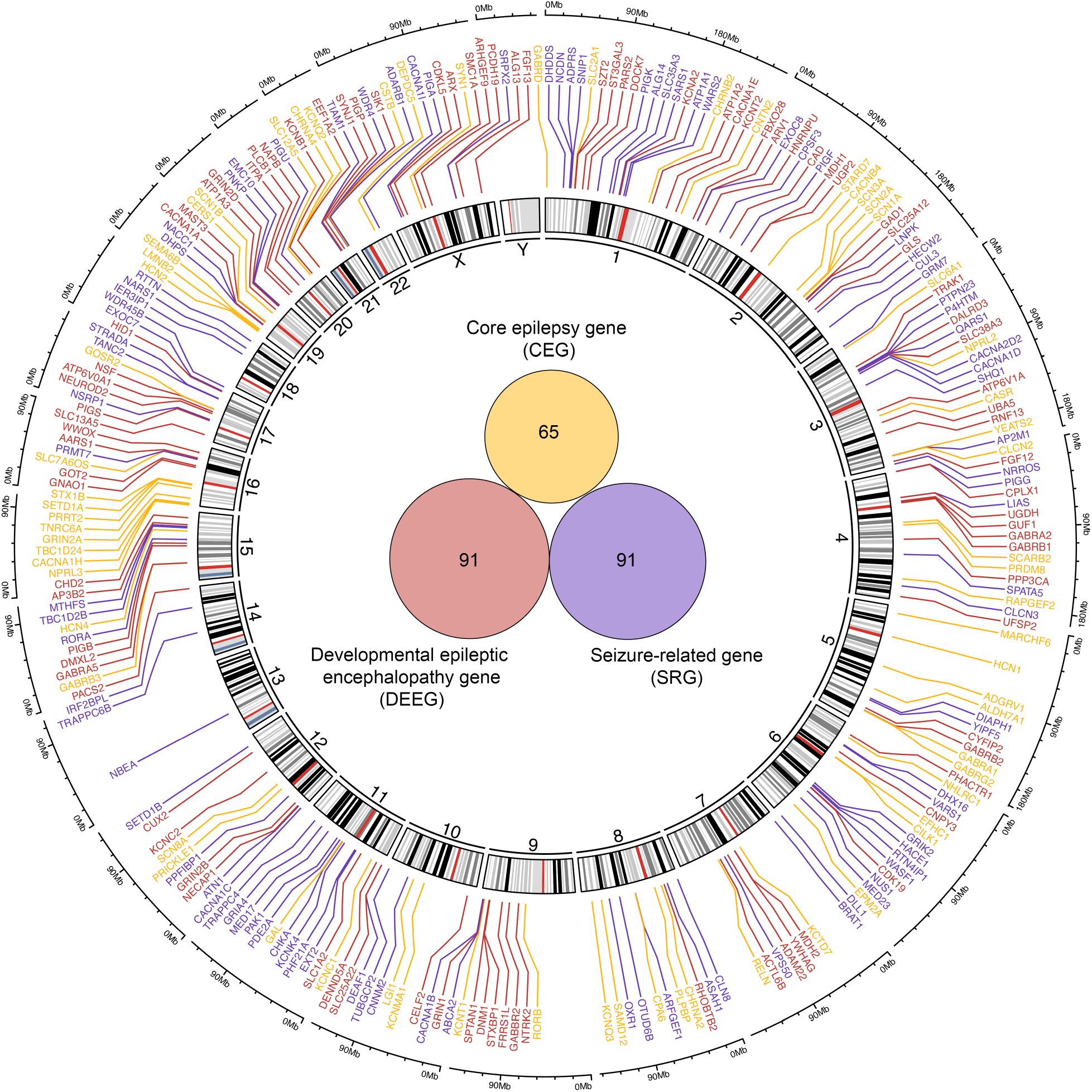
Genomic profile of epilepsy-associated genes. A Circos plot representing the genomic location of epilepsy-associated genes. Epilepsy-associated genes that are color-coded based on phenotypes identified in patients carrying mutations in corresponding genes. Inside the Circos is a Venn diagram that shows the number of genes in each group. Genes and associated phenotypes are obtained from OMIM.

Epilepsy-associated genes are encoded relatively uniformly across the genome (Fig. 1). Each chromosome contains multiple genes except for chromosomes 13 and Y that have one and none respectively. Notably, epilepsy-associated genes are enriched for various functional categories including “enzyme” (27.5%), “ion channel” (16.6%), “transporter/receptor” (11.7%), “signaling” (14.2%), “transcription/translation” (10.5%), “cytoskeleton” (2.8%), and “extracellular matrix” (0.8%), while several genes remain uncharacterized (10.9%) (Fig. 2). The diverse protein functions of epilepsy-associated genes suggest that neuronal activity in the brain is intricately modulated.

**Figure 2.**
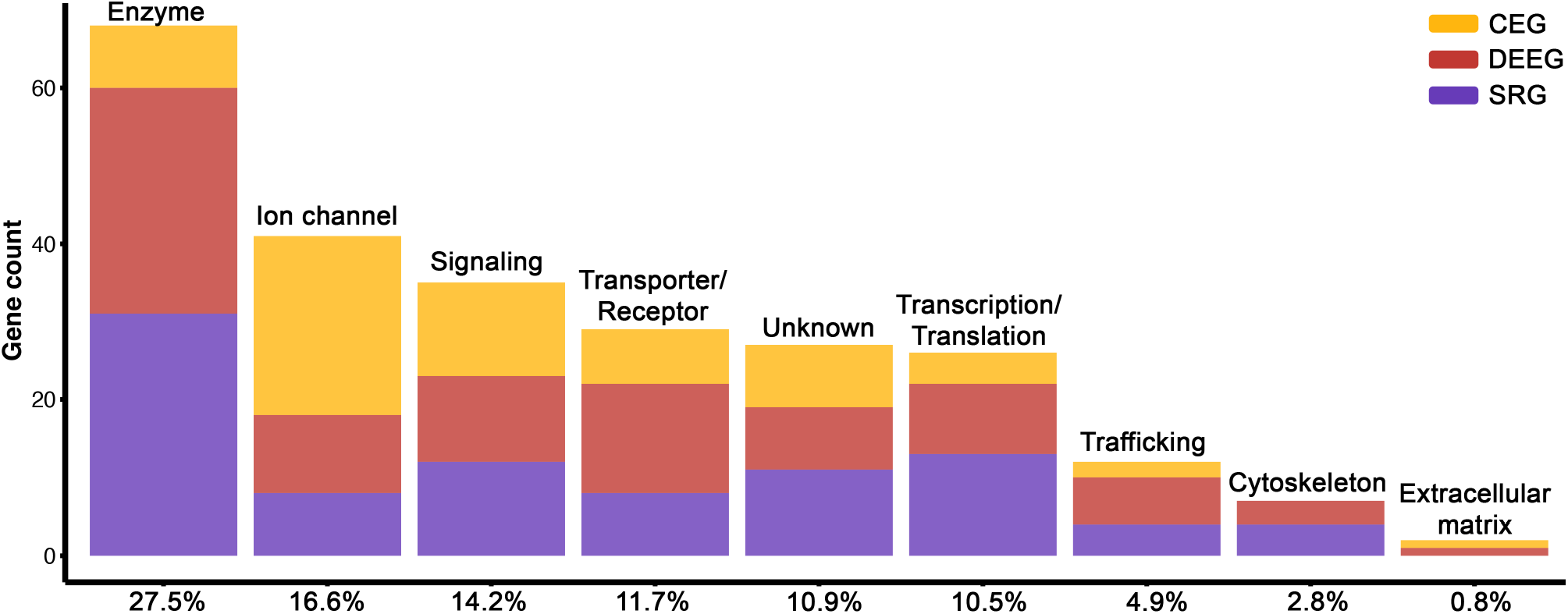
Functional classification of epilepsy-associated genes. Protein Class categories for 247 epilepsy-associated genes based on PANTHER. The percent of each category relative to all genes is shown on the X-axis.

### The three groups of epilepsy-associated genes exhibit differential expression between CNS and non-CNS tissues

To assess the pattern of expression for epilepsy-associated genes, we first obtained tissue-specific expression data from GTEx^7^, which includes 14 CNS tissues and 38 non-CNS regions. We performed hierarchical clustering analyses across all genes and samples and found a clear separation between brain regions (and testis) and non-CNS tissues (Fig. 3A). In terms of epilepsy gene groups, DEEGs showed the highest expression within the CNS, followed by CEGs and SRGs which on average were comparable (Fig. 3B-D). In contrast, genes within the SRG group were most highly expressed in non-CNS tissues, followed by DEEGs and CEGs. The elevated expression of DEEGs in the brain may explain the reduced incidence of peripheral tissue comorbidities manifested by patients carrying such mutations, while the more ubiquitous expression of SRGs correlates with the higher incidence of diverse comorbidities found in SRG-associated patients. Individual genes identified as major contributors to epilepsy such as *SCN1A, SCN2A* and *KCNQ2* exhibited diverse expression within regions of the brain (Supplemental Fig. 1).

**Figure 3.**
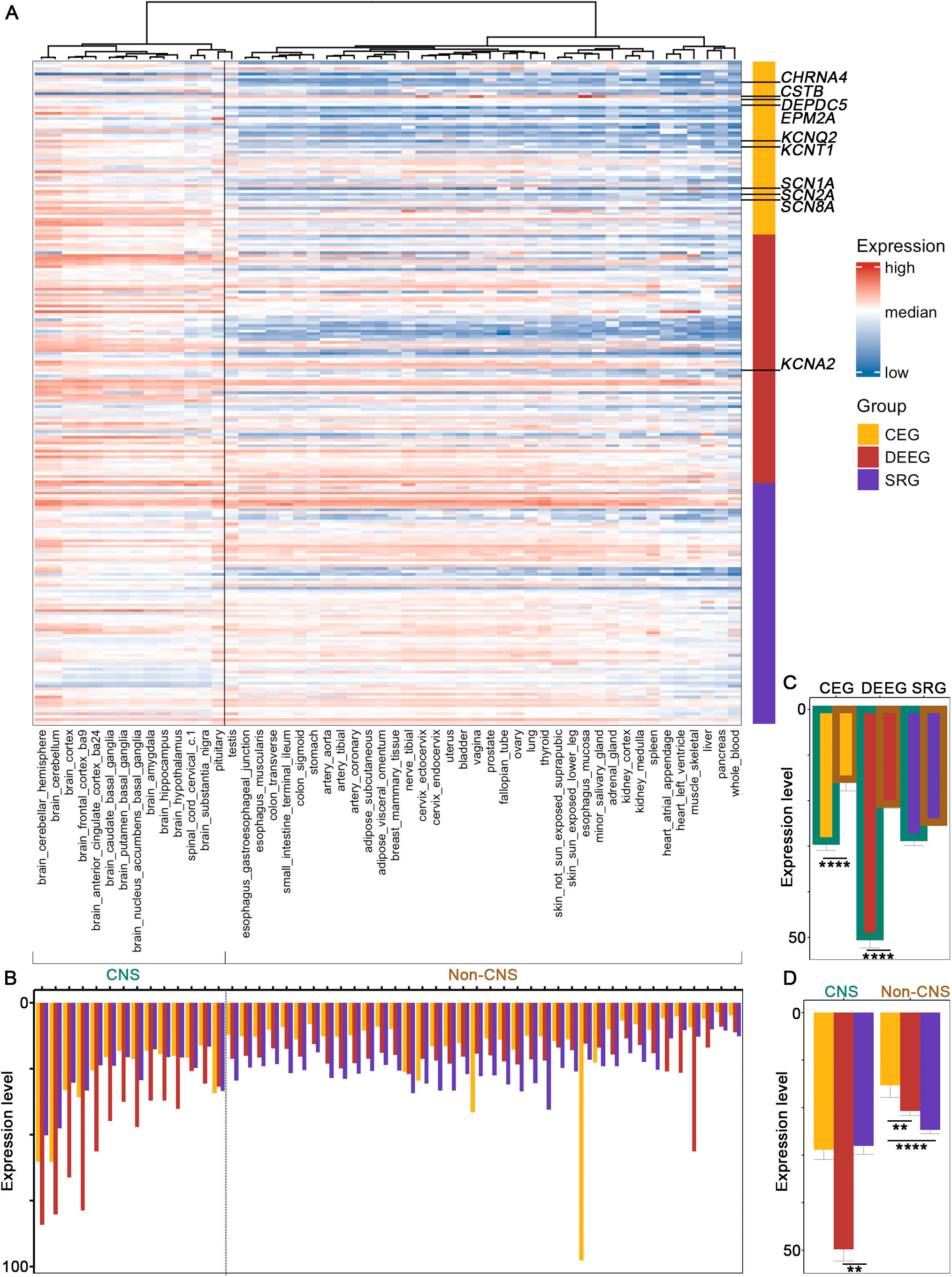
Expression of the three groups of epilepsy-associated genes in human tissues. (A) Heatmap for hierarchical clustering. Top 10 studied genes^6^ are labelled (see Supplemental Fig 1 for individual plots). (B) Averaged expression of all the genes in each group across different tissues. (C,D) Averaged expression of all the genes in each group in CNS versus non-CNS tissues. ** p < 0.01, ****p < 0.0001, Wilcoxon signed rank sum test with Benjamini-Hochberg post hoc. Bulk RNA-seq expression data are from GTEx.

### DEEGs exhibit high expression across development

Given that DEE patients exhibit neurodevelopmental deficits, we next examined the developmental expression pattern of epilepsy-associated genes in the brain. We obtained RNA-seq data from different stages of human development, ranging from 4 post conception weeks (PCW) to 60+ years, in 26 brain sub-regions generated by the Allen Brain Institute^9^ (Fig. 4A-B). For clarity, we grouped the datasets into one of five developmental stages: prenatal, infancy, childhood, adolescence, and adulthood (Fig. 4A). As a group, DEEGs exhibited significantly higher expression levels (2-3-fold) relative to CEGs and SRGs within the hippocampus, as well as greater variation during development, rising from prenatal stages to infancy, normalizing during childhood, and then slowly increasing again in adolescence and adulthood (Fig. 4C-D; Supplemental Fig. 2). Close examination of individual gene expression across the five developmental periods within the hippocampus revealed that on average both DEEGs and CEGs transition into higher expression levels during the shift from prenatal to infancy, while SRGs were the least dynamic (Fig. 4D). These patterns were similarly observed across the other 15 brain regions we examined with minor variations (Supplemental Fig. 3). Interestingly, although the average expression of CEGs is comparable to SRGs in the prenatal period, it becomes consistently higher than SRGs in later stages of development, such that during adulthood expression of CEGs is significantly higher relative to SRGs in 13 out of 16 studied regions (Supplemental Fig. 4).

**Figure 4.**
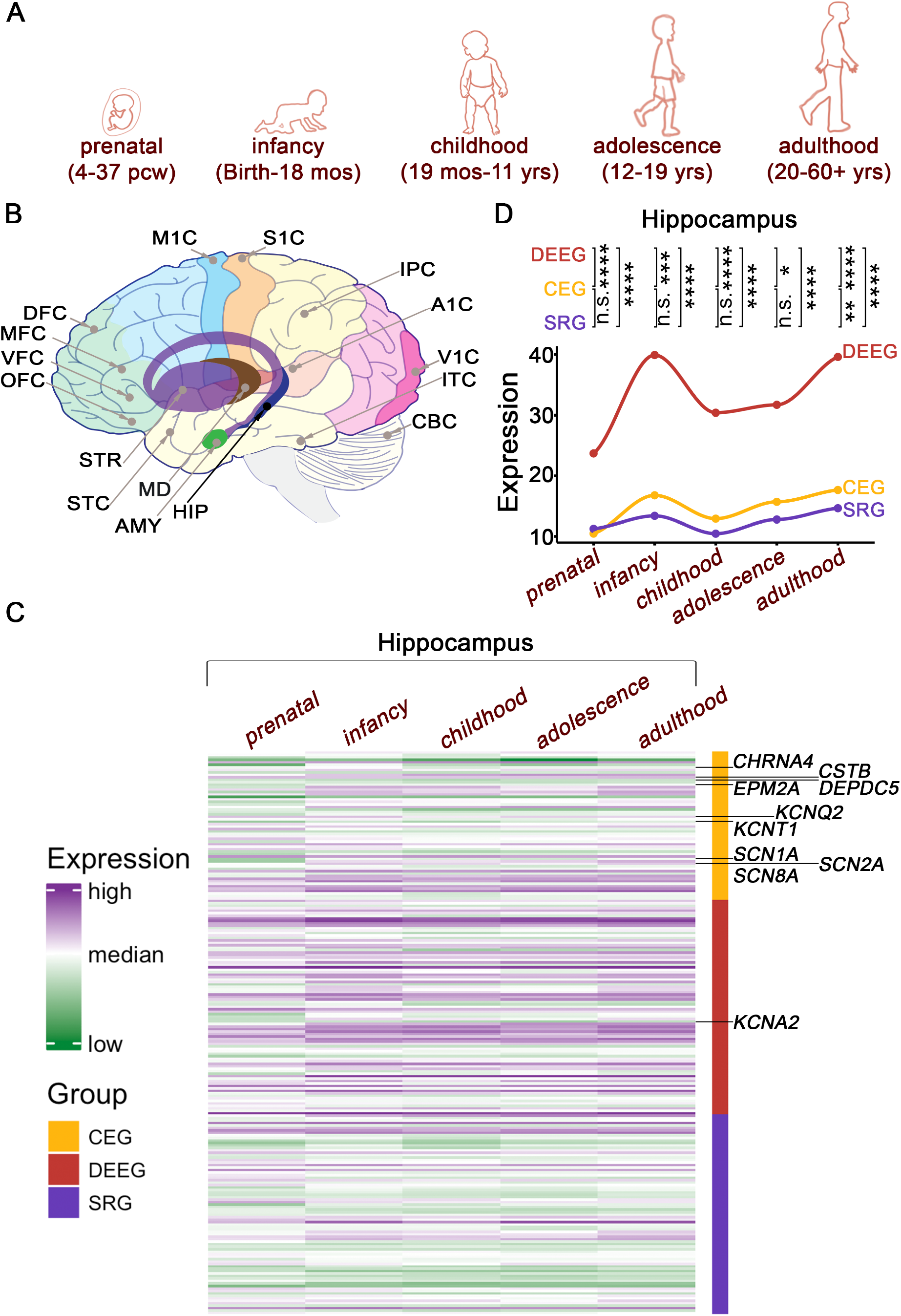
Hierarchical clustering of epilepsy-associated genes in the developing brain. (A) A cartoon presentation of human development with respective ages. (B) Brain sub-regions included in the analyses. A1C: primary auditory cortex; AMY: amygdala; CBC: cerebellar cortex; DFC: dorsolateral prefrontal cortex; HIP: hippocampus; IPC: posteroventral parietal cortex; ITC: inferolateral temporal cortex; M1C: primary motor cortex; MD: Mediodorsal nucleus of thalamus; MFC: anterior cingulate cortex; OFC: orbital frontal cortex; S1C: primary somatosensory cortex; STC: posterior superior temporal cortex; STR: striatum; V1C: primary visual cortex; VFC: ventrolateral prefrontal cortex. (C) Heatmap from hippocampus. Top 10 studied genes^6^ are labelled (see Supplemental Fig 2 for individual plots). (D) Averaged expression of all the genes within each group across development in the hippocampus. n.s. p > 0.05, ** p < 0.01, ****p < 0.0001, Wilcoxon signed rank sum test with Benjamini-Hochberg post hoc. See Supplemental Figs 3,4 for other regions. Bulk RNA-seq data are from Allen BrainSpan.

### Expression of epilepsy-associated genes in neurons does not correlate with phenotypes

We further examined the expression of the three groups of epilepsy-associated genes in brain-specific cell types. We utilized an extensive set of single-nucleus human RNA-seq data from the Allen Brain Institute^8^, which contains sets from GABAergic and glutamatergic neurons and 4 types of non-neuronal cells: astrocytes, oligodendrocyte progenitor cells (OPCs), oligodendrocytes, and microglia. Within each neuronal cell type, individual cells are further classified into 24 subtypes of glutamatergic neurons and 17 subtypes of GABAergic interneurons based on the expression of specific molecular markers.

Hierarchical clustering analysis based on the expression of all 247 epilepsy genes across brain subtypes revealed a clear separation into three distinct groups: glutamatergic neurons, GABAergic interneurons, and non-neuronal cells (Fig. 5A). While most epilepsy genes showed relatively similar expression between GABAergic and glutamatergic neurons, some genes such as *GAD1* and *SLC6A1* that are cell type-specific markers, were significantly different between these two neuronal cell types (Supplemental Fig 5). Lastly, our analysis revealed that the average expression of CEGs and SRGs within cellular subtypes was largely comparable (Fig. 5B-C), while the average expression level of DEEGs was significantly higher than the other two groups in GABAergic neurons and non-neuronal cells (Fig. 5C).

**Figure 5.**
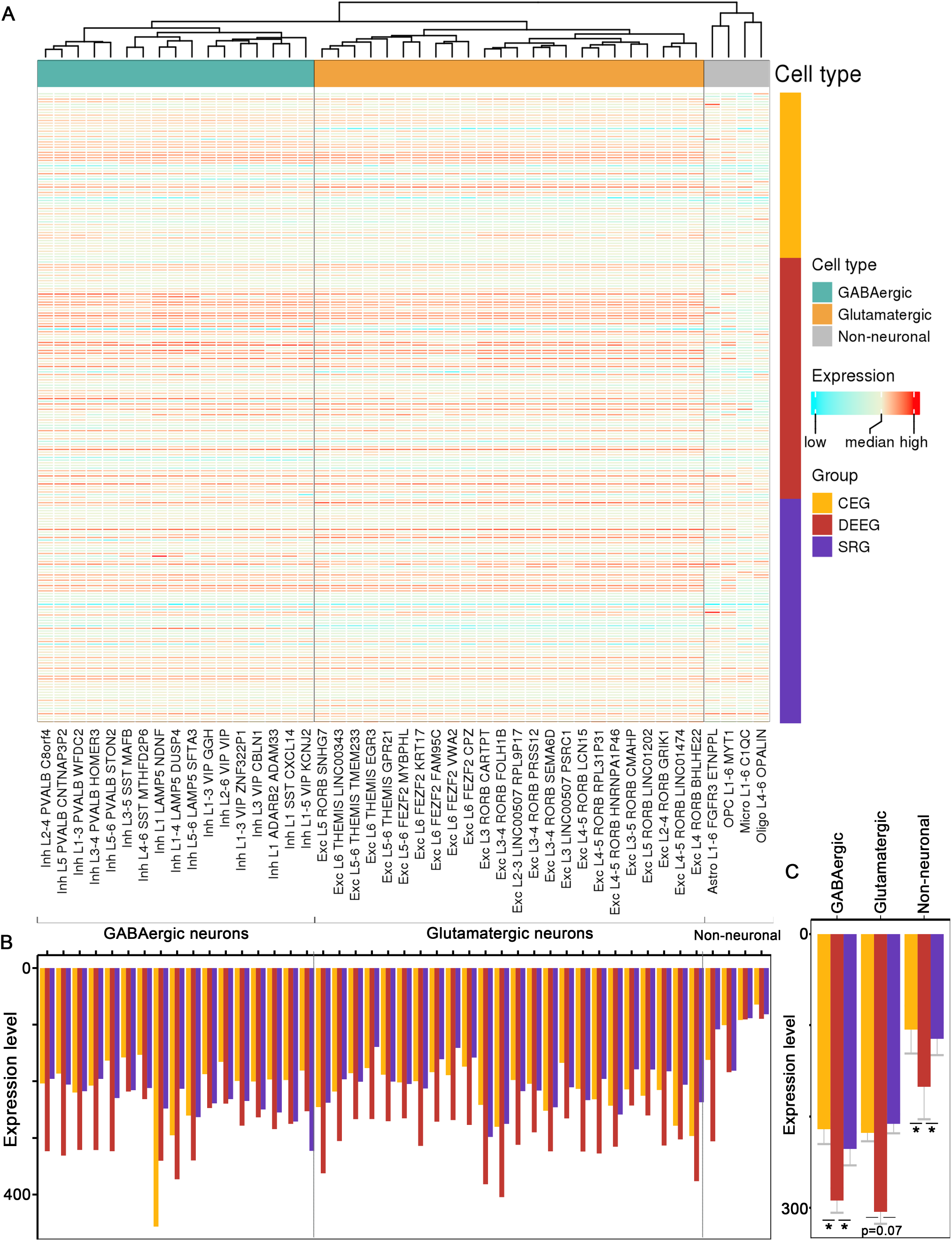
Expression of epilepsy-associated genes in different brain cell types. (A) Heatmap for hierarchical clustering across cell types. see Supplemental Fig 5 for individual plots of top 10 studied genes^6^. (B) Averaged expression of all the genes within each group across different cell type clusters. (C) Averaged expression of all genes in each group across three cell type classes. * p < 0.05, Wilcoxon signed rank sum test with Benjamini-Hochberg post hoc. Single-nucleus RNA-seq data of each cell type are from Allen BrainMap.

## Discussion

Here we systematically reviewed the expression pattern of 247 epilepsy-associated genes in human tissues, across development, and within brain-specific cell types to explore the relationship between gene expression and phenotypic manifestation.

Epilepsy is a common neurological disorder characterized by phenotypic diversity with a substantial genetic component to its etiology. Based on the symptoms curated in OMIM, we grouped epilepsy-associated genes into CEGs, DEEGs, and SRGs. Patients with CEGs manifest seizures and rarely have comorbidities, patients with DEEGs show seizures and neurodevelopmental deficits attributed to epileptic activity, and patients with SRGs exhibit seizures and other comorbidities as well as structural deficits such as brain malformation. Our analysis demonstrates that DEEGs have the highest expression in the brain across development and that SRGs have the highest expression in non-CNS tissues. The high expression levels of genes within these groups indicate their important physiological roles in respective tissues, which, when disrupted by disease-associated mutations, likely contribute to the various comorbidities associated with these conditions.

Epilepsy has been postulated to arise as a result of a network imbalance^14^, likely involving both excitatory and inhibitory neurons. Our cell type analyses demonstrates that most epilepsy-associated genes are similarly expressed in both types of neurons at least in cortical brain regions. At the same time, DEEGs as a group were significantly more abundant in interneurons and non-neuronal cells, suggesting a potential contribution of inhibitory signaling and oligodendrocytes and glial cells in these epilepsy cases.

Importantly, we found that some genes are associated with divergent clinical phenotypes. For example, *SCN1A, SCN2A, SCN3A, SCN8A, SCN1B, SLC12A5, TBC1D24, GABRA1, GABRB3, GABRG2, HCN1, KCNQ2*, and *KCNT1* are shared between the CEG and DEEG groups. While we included these genes within the CEG group in our analyses, the fact that they are involved in two phenotypic outcomes suggests that the gene expression pattern cannot be the only source of phenotypic diversity. As a matter of fact, many studies have demonstrated that several factors can contribute to phenotypic diversity. First, the functional effect of the mutation on the encoded protein matters, which is especially true for ion channels. One such prominent example is the *SCN1A* gene. While loss-of-function (LOF) variants in *SCN1A* gene cause Dravet syndrome and genetic epilepsy with febrile seizures plus, gain-of-function (GOF) variants are associated with familial hemiplegic migraine^15,16^. Mechanistically, LOF and GOF *SCN1A* mutations can lead to either enhanced or diminished *SCN1A* ion channel activity, which will lead to differential neuronal excitability effects depending on the neuronal cell type. Consistent with this notion, single cell expression analyses show that *SCN1A* is expressed similarly in GABAergic and glutamatergic neurons. In addition, Thompson and colleagues recently showed that different *SCN2A* variants exert different functional effects on neonatal versus adult *SCN2A* isoforms, leading to complex phenotypic outcomes in immature versus mature neurons^17^.

Second, diverse phenotypes could also arise as a result of the interaction between the gene and other genetic and environmental factors. Gene-gene interactions have been shown for several ion channels including *SCN1A, SCN2A, SCN8A, KCNQ2, KCNA1, CACNA1G* and *CACNA1A*^18–23^. Notably, *SCN1A, SCN2A, SCN8A*, and *KCNQ2* are shared by CEG and DEEG in our analyses. Lastly, genetic mosaicism has been increasingly recognized to contribute to phenotypic diversity in human diseases including epilepsy^24^. Thus, while collectively there is a correlation between gene expression and phenotypic presentation, each individual gene itself also compasses complexity due to the mutation properties, gene-gene, and gene-environment interactions, as well as genetic mosaicism.

Limitations of our work include the fact that we only analyzed 247 epilepsy-associated genes and grouped them solely based on the clinical phenotypes curated in OMIM. While this database provides accurate gene-phenotype associations, the list of genes is continuously refined based on the latest data. For instance, many new genes have been identified but have yet to be curated in the database, while three recent studies^5,6,25^ show that the number of epilepsy-associated genes is close to 1000, although the exact number can vary based to the inclusion and exclusion criteria applied. Xian and Helbig recently reported that *CACNA1H, PRICKLE1, CACNB4, EFHC1, MAGI2, SCN9A, CASR, SRPX2* should no longer be considered epilepsy genes because of the lack of definitive variant-disease relationship^26^. Additionally, many reported genetic variants remain functionally undefined and thus their contribution to disease pathophysiology is unknown. Lastly, the clinical phenotypes associated with epileptic syndromes are often complex and lack of standardized descriptions may confound the precision of gene-phenotype curation in OMIM.

Despite these limitations, our study provides an overview of the expression pattern of an inclusive collection of epilepsy associated genes across tissues, developmental stages, and neural sub-types and can serve as a useful resource for the community.

## Acknowledgments

We thank Dr. Nan Xiao for technical assistance.

## Author contribution

Resource design and conceptualization: W.C., E.K. Analysis and Drafting the manuscript: W.C. All authors read and approved the final version of the manuscript.

