## Supplementary figures and images for "Integrative Analysis of Epilepsy-Associated Genes Reveals Expression-Phenotype Correlations"

### Supplemental Fig 1

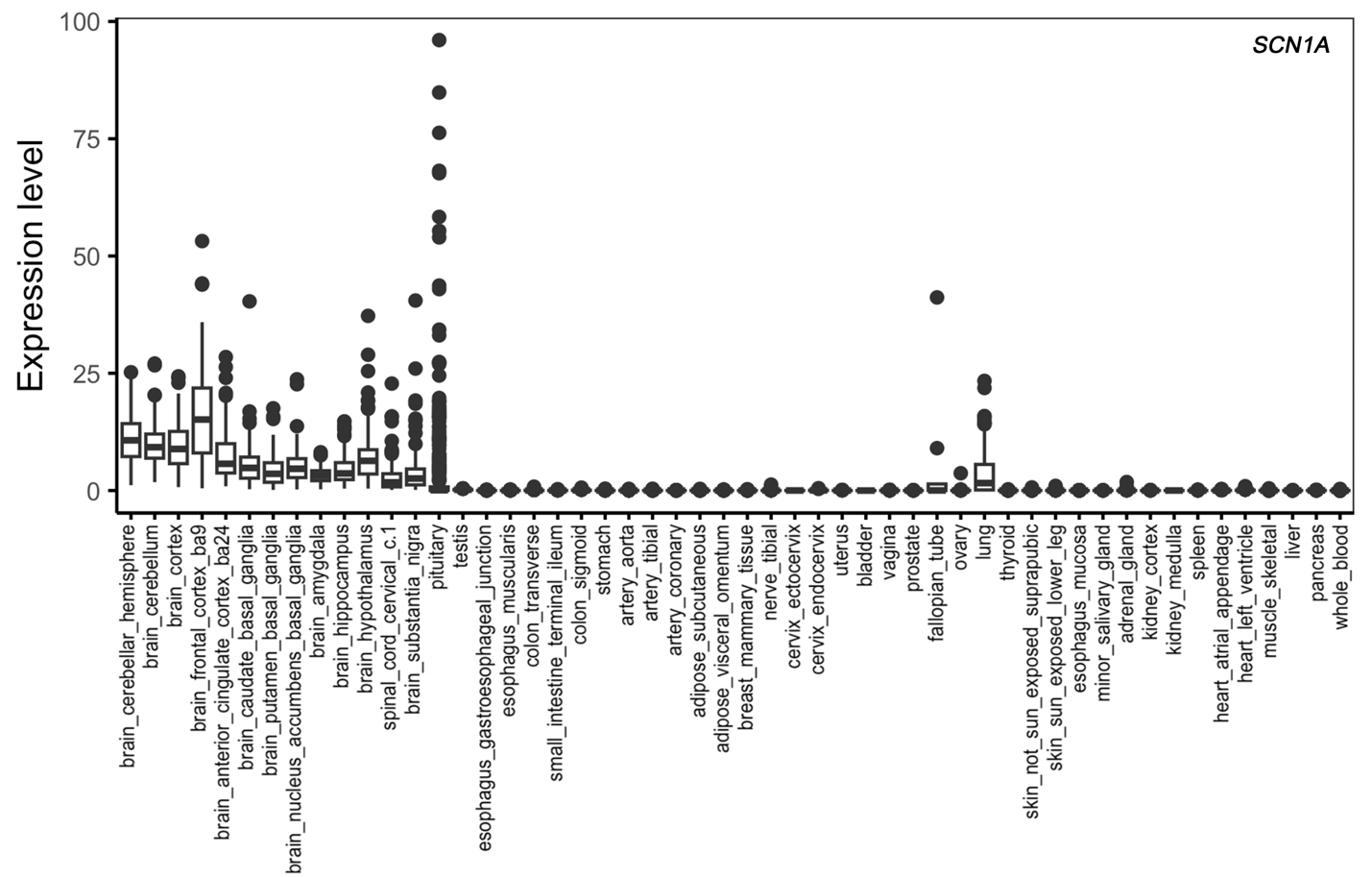

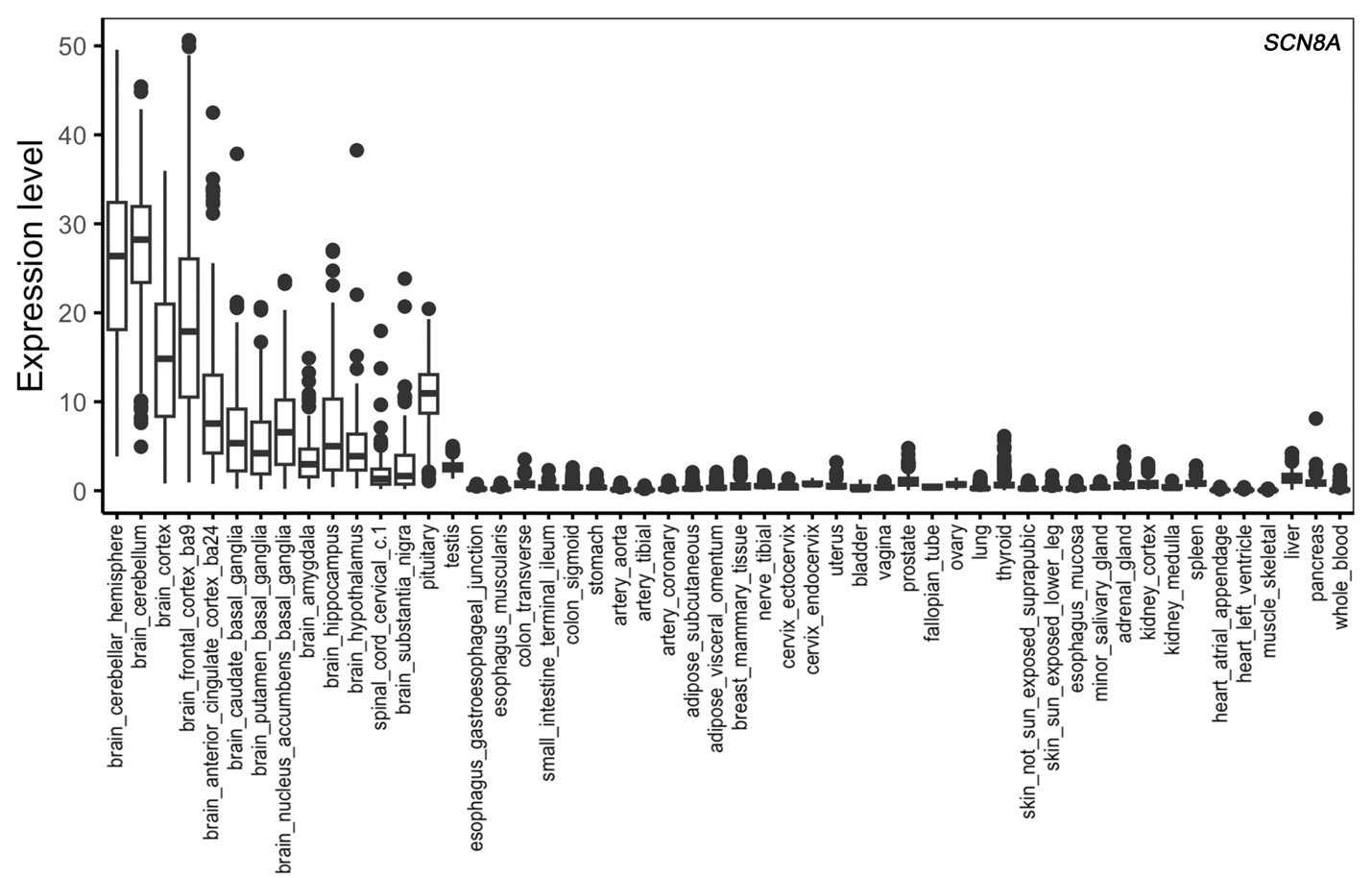

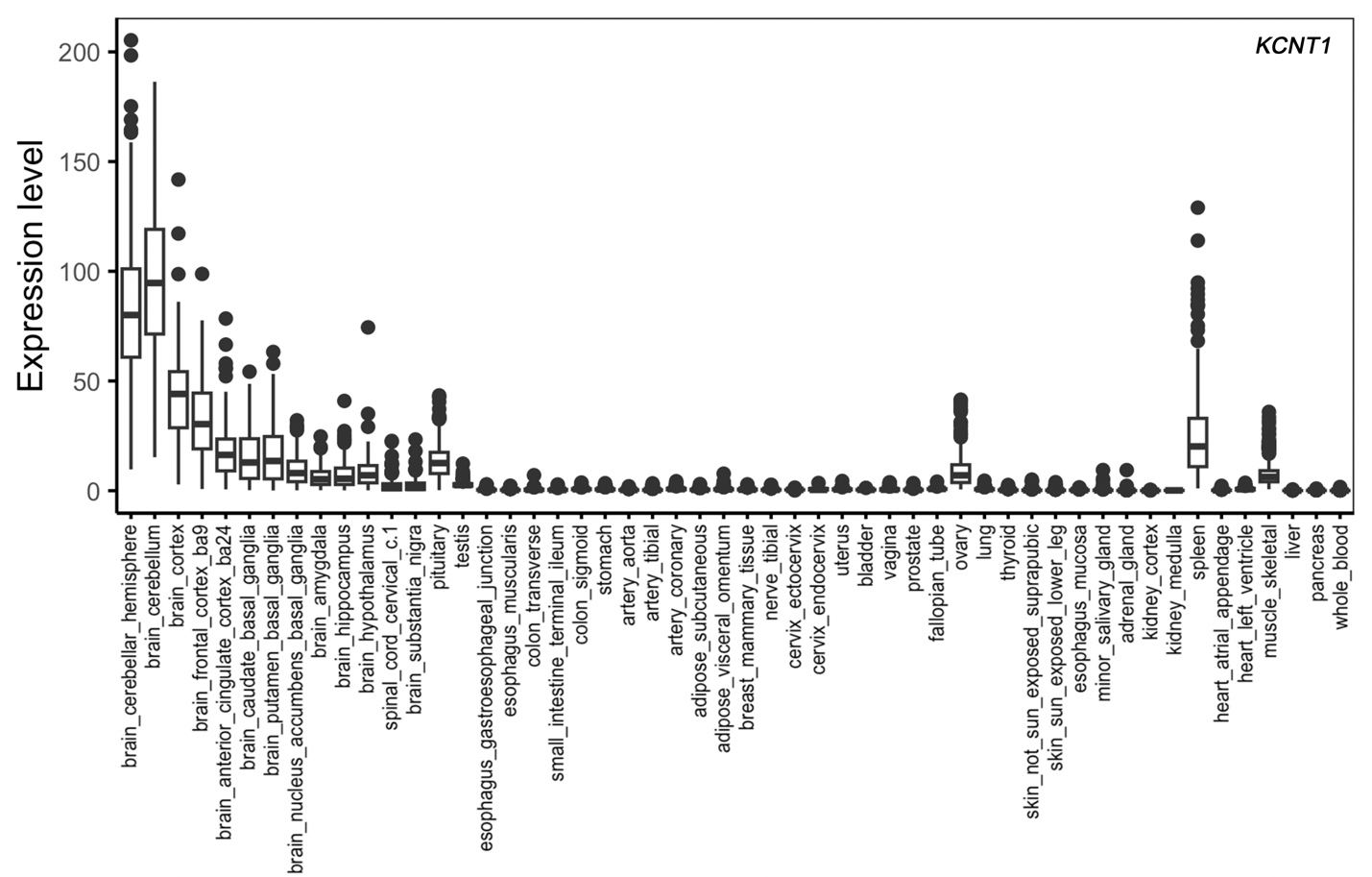

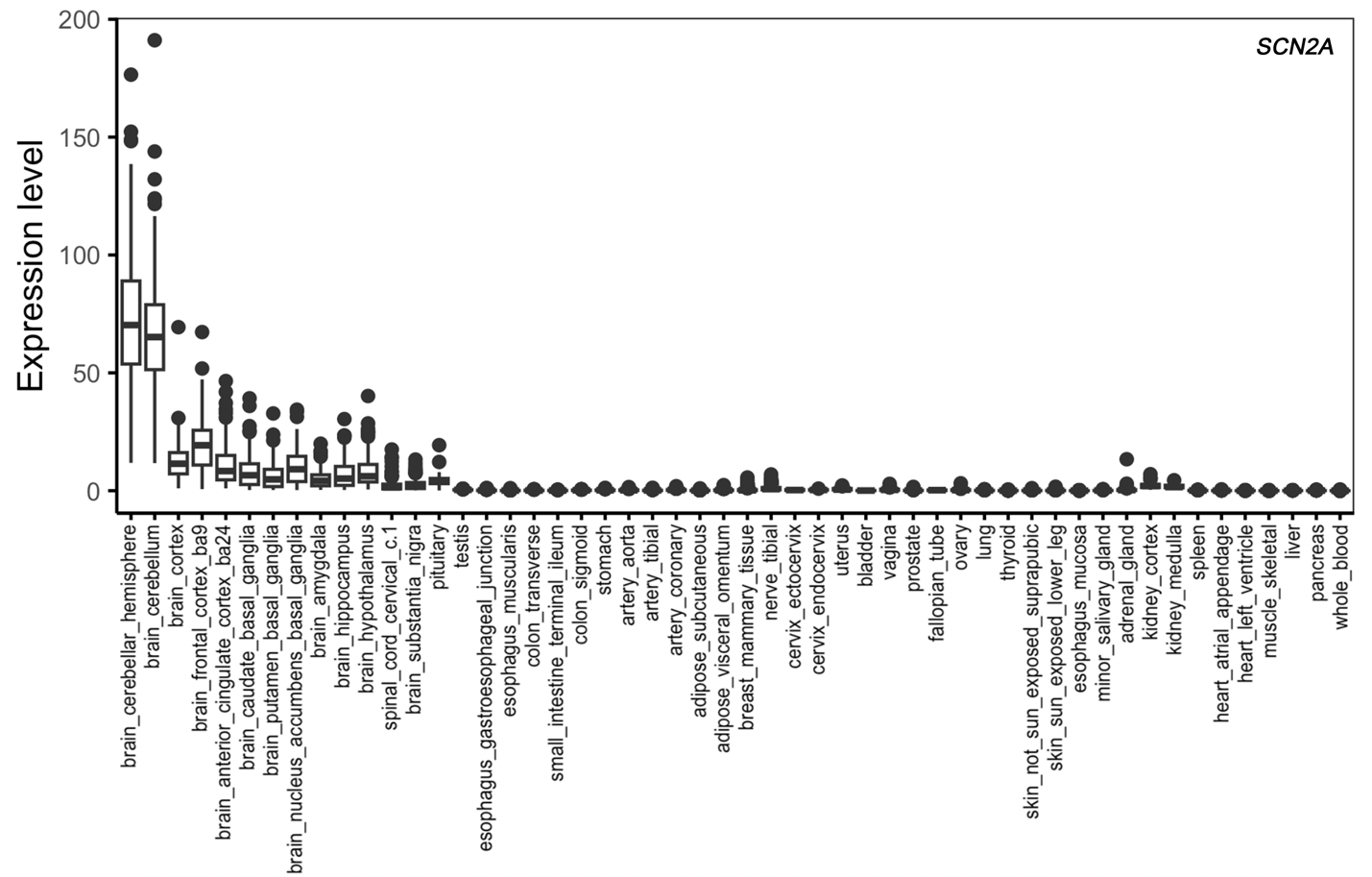

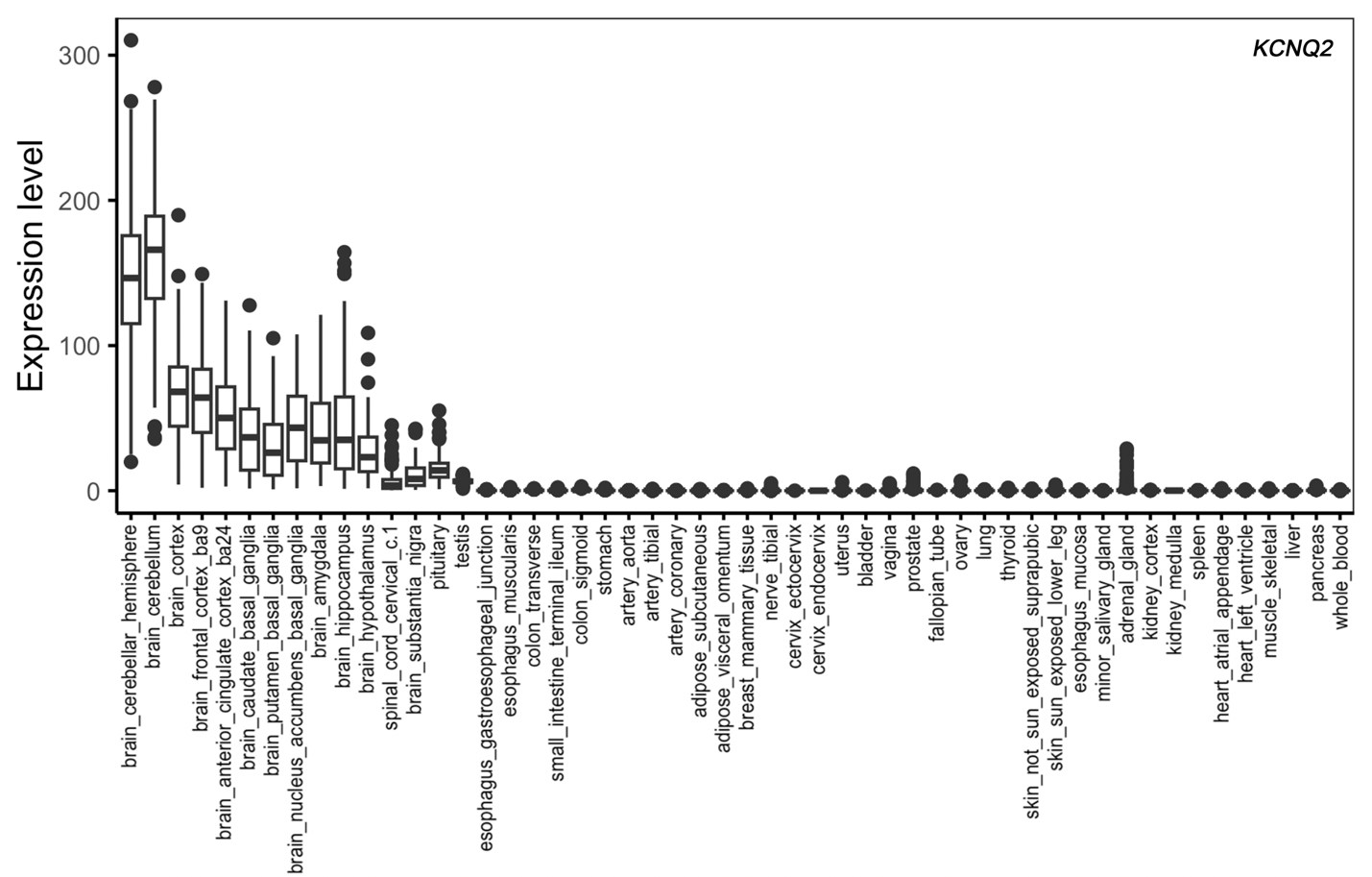

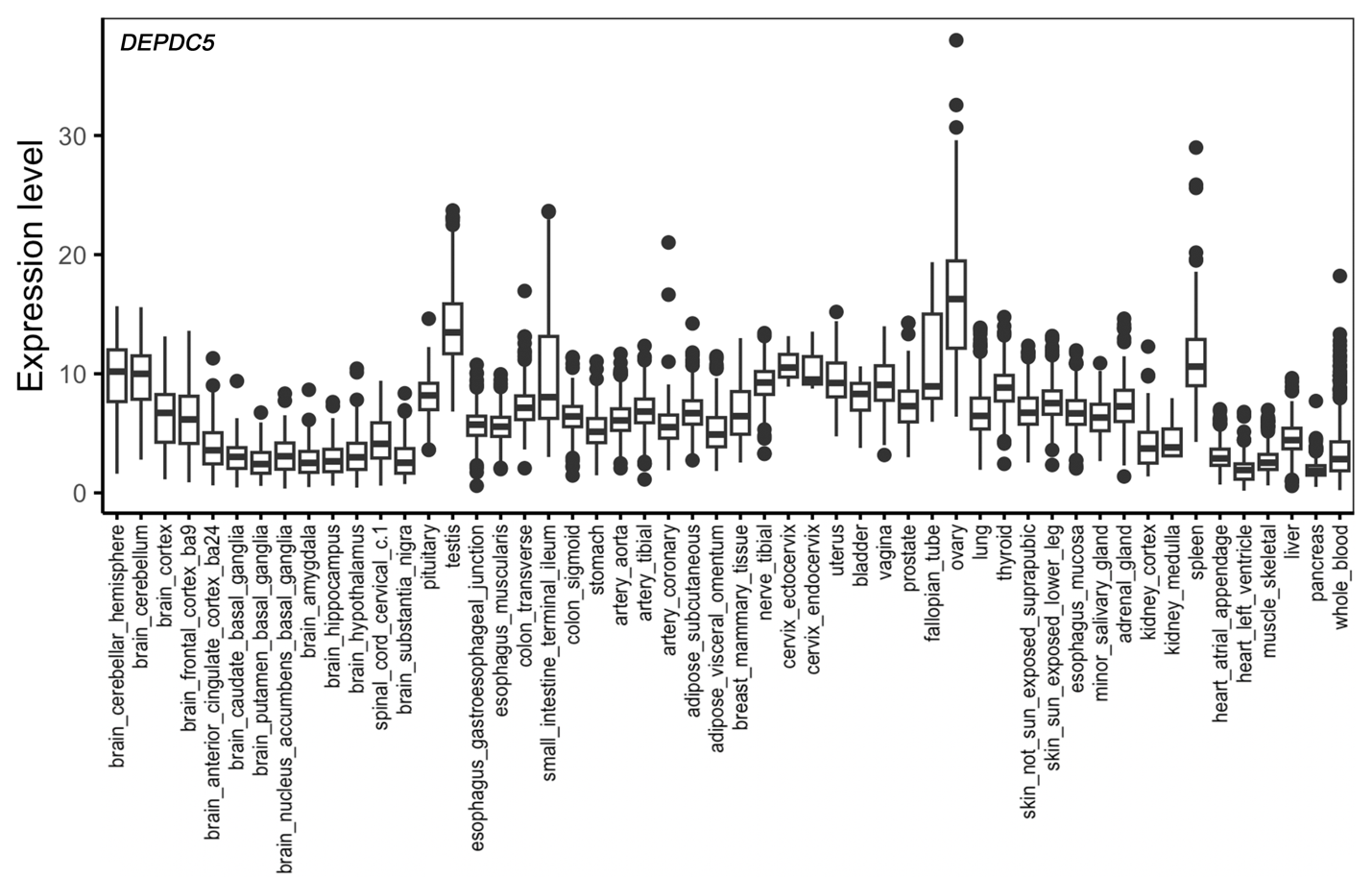

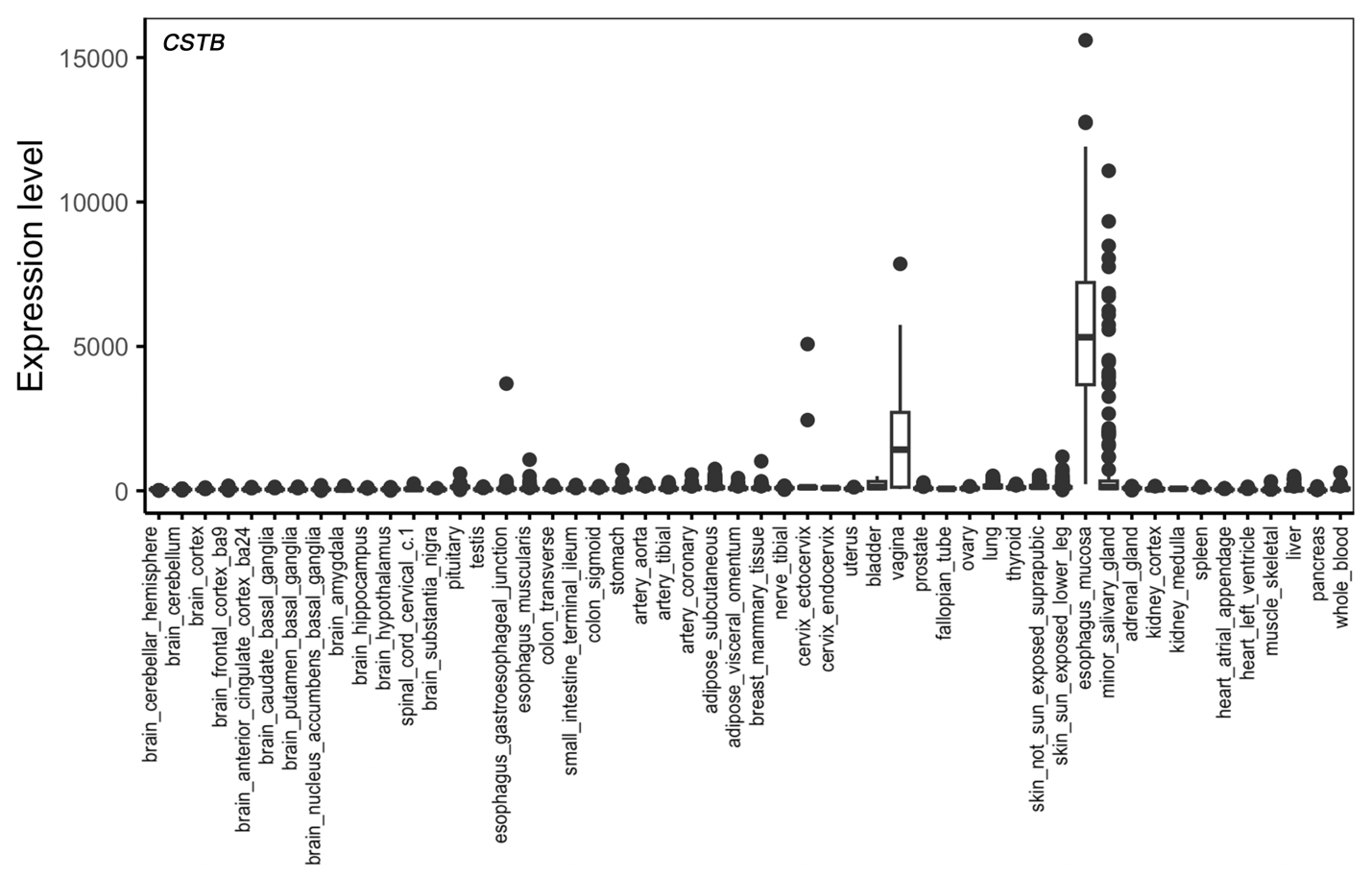

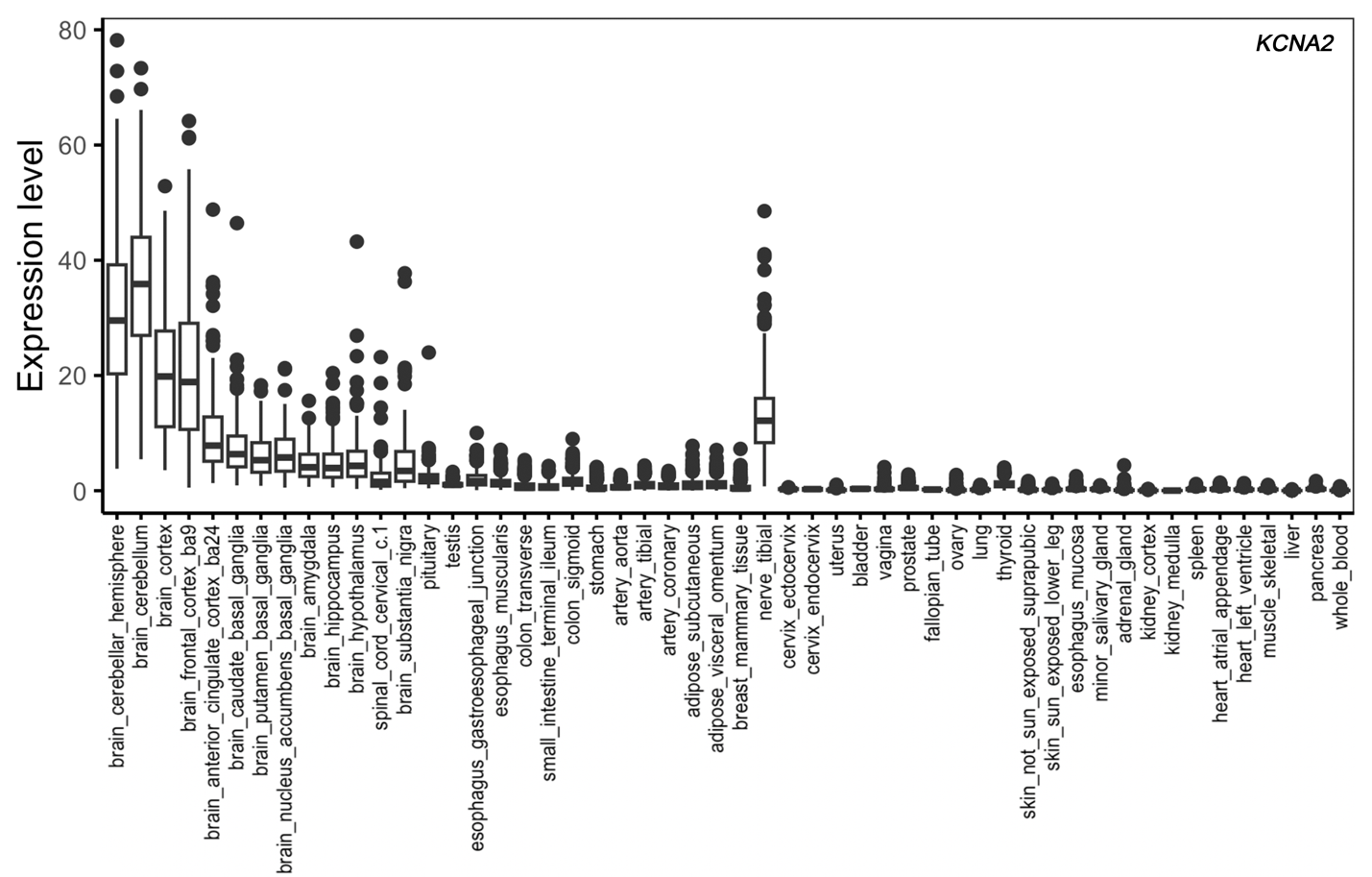

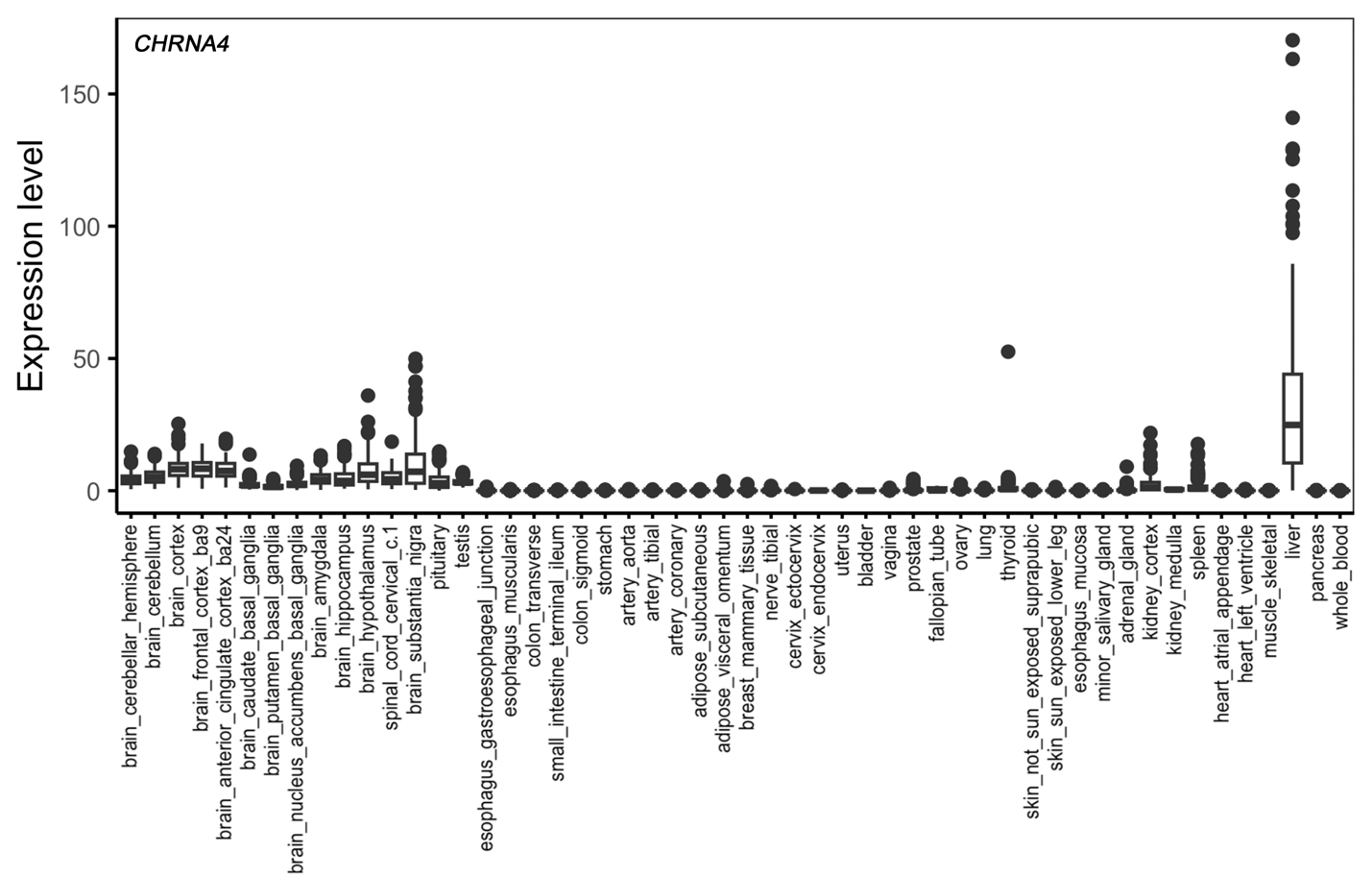

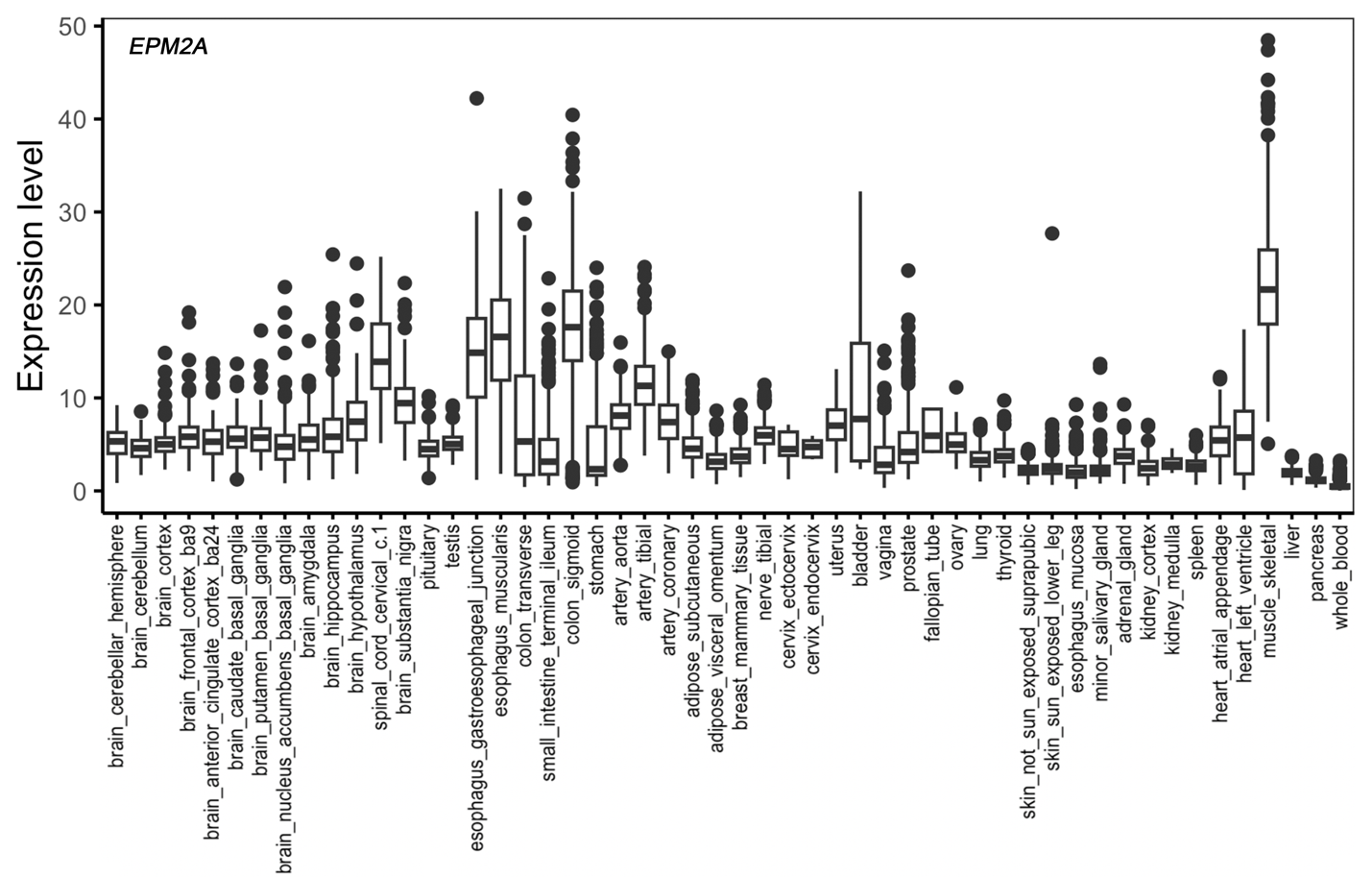

**Supplemental Figure 1. Expression of top 10 studied genes across human tissue.**
