## Supplemental Fig 2 for "Integrative Analysis of Epilepsy-Associated Genes Reveals Expression-Phenotype Correlations"

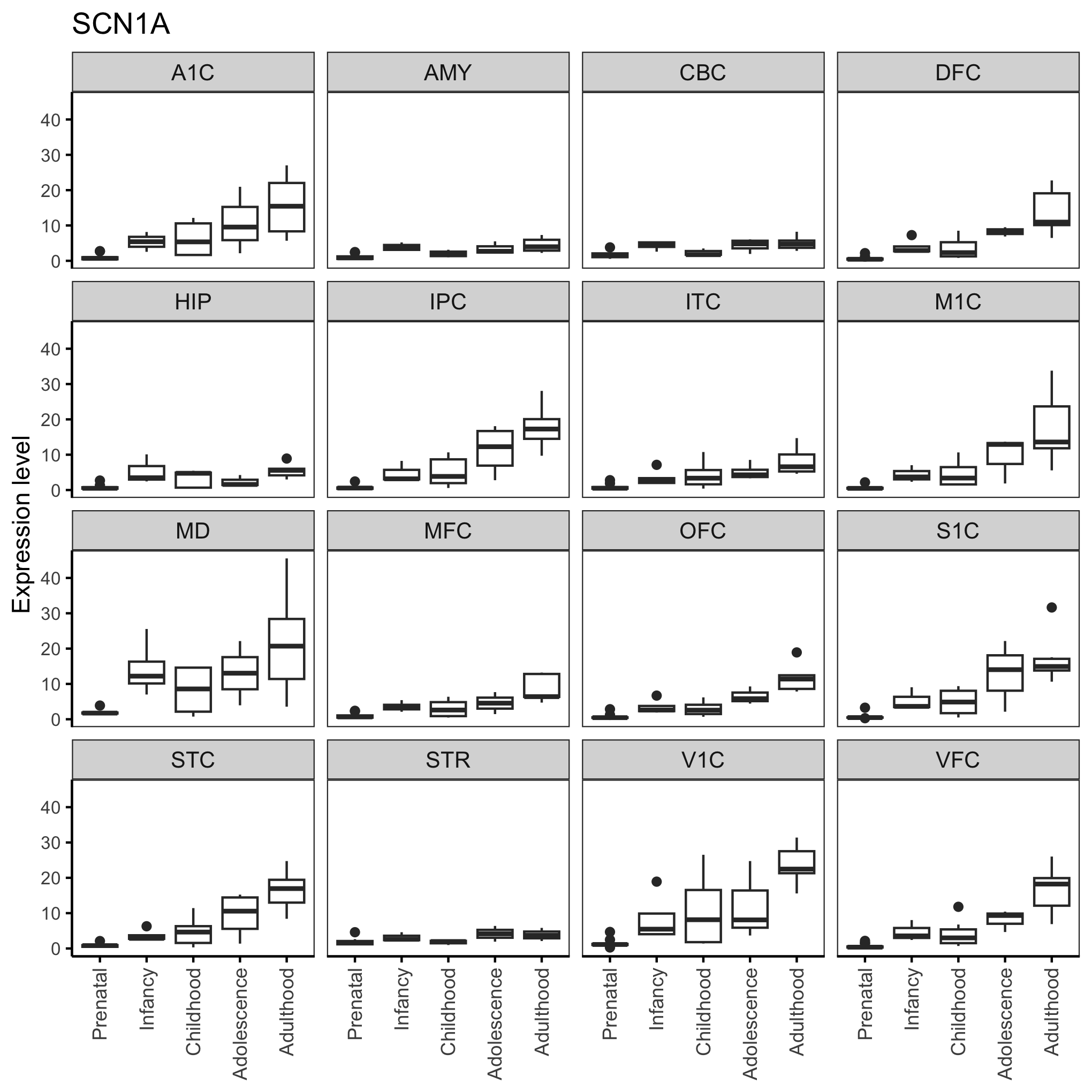

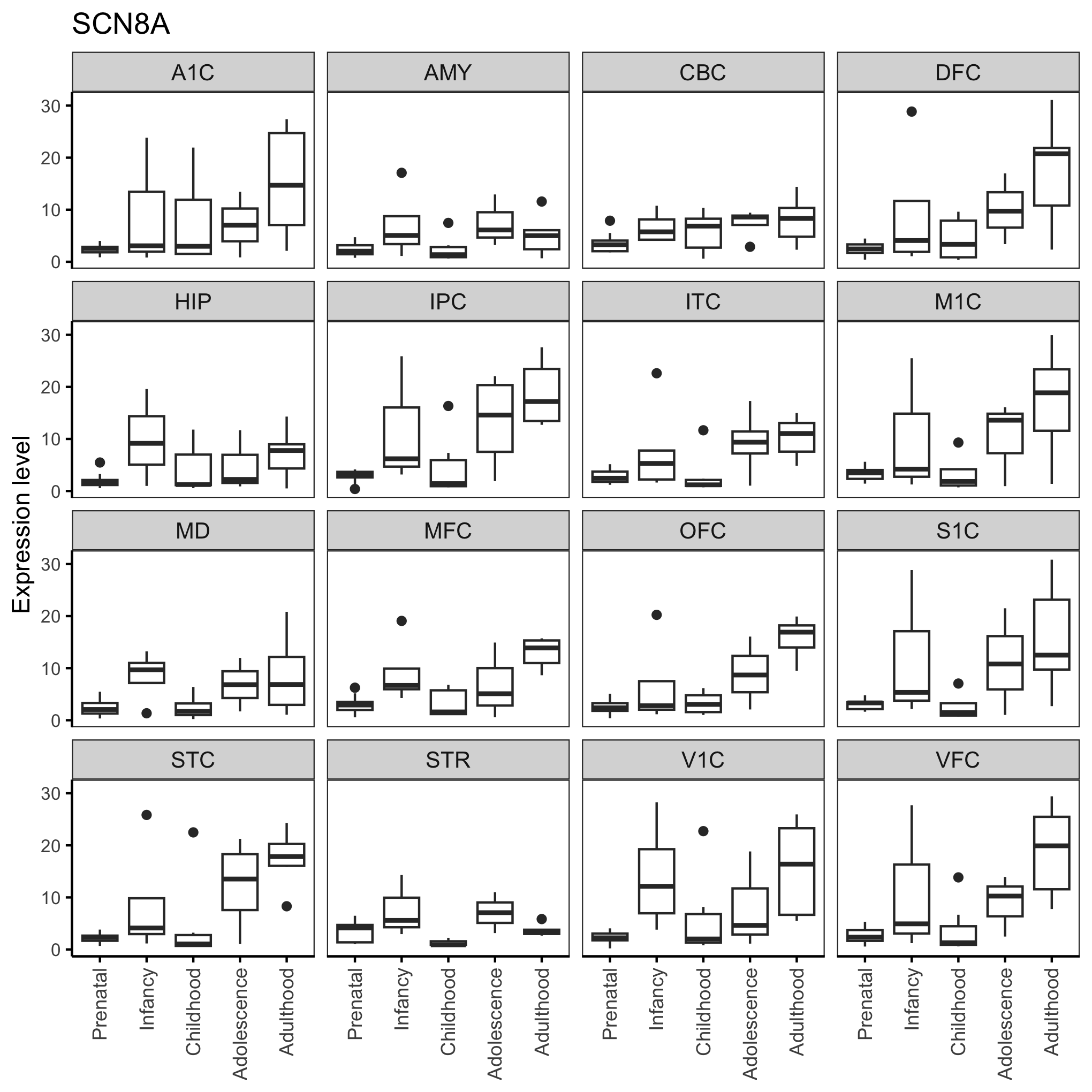

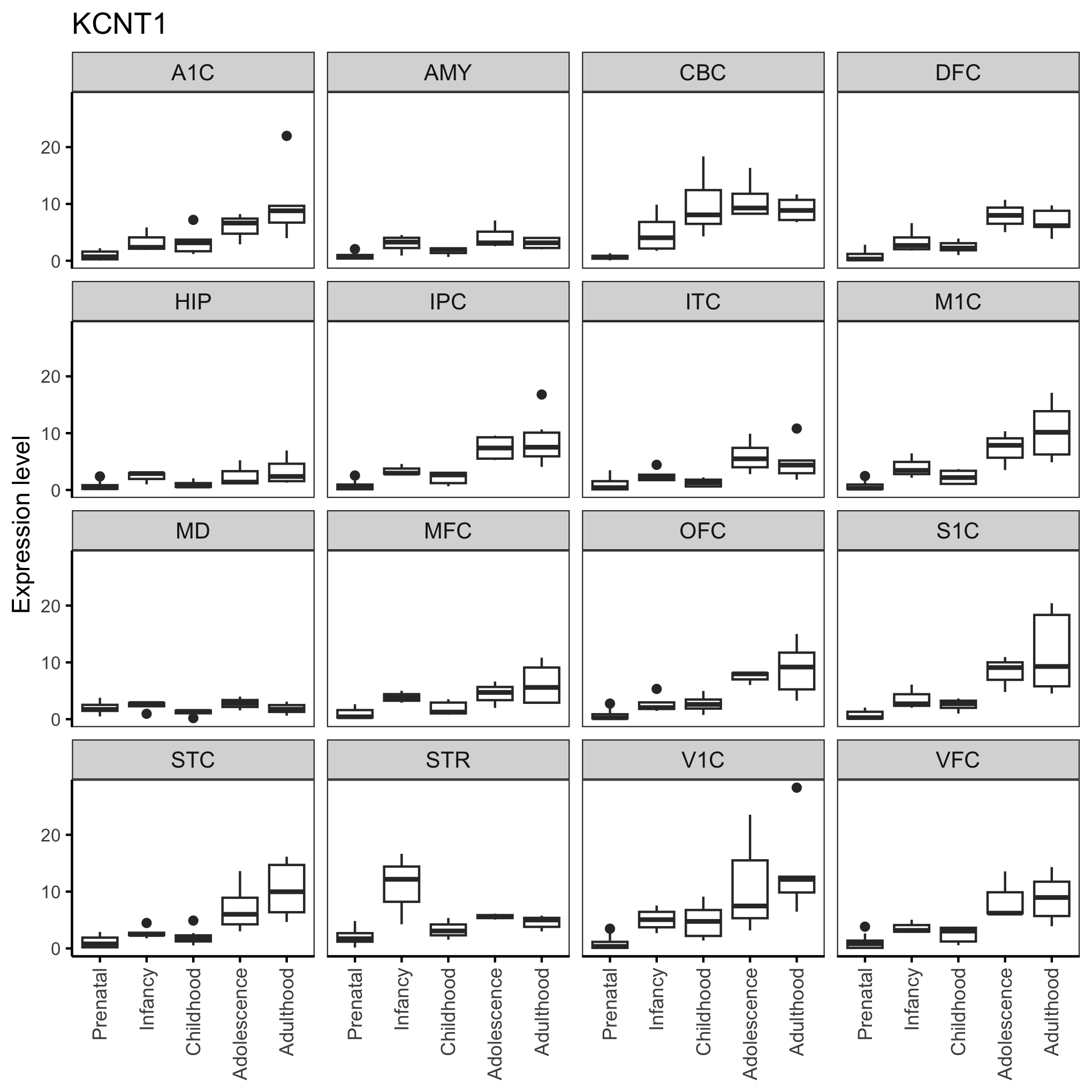

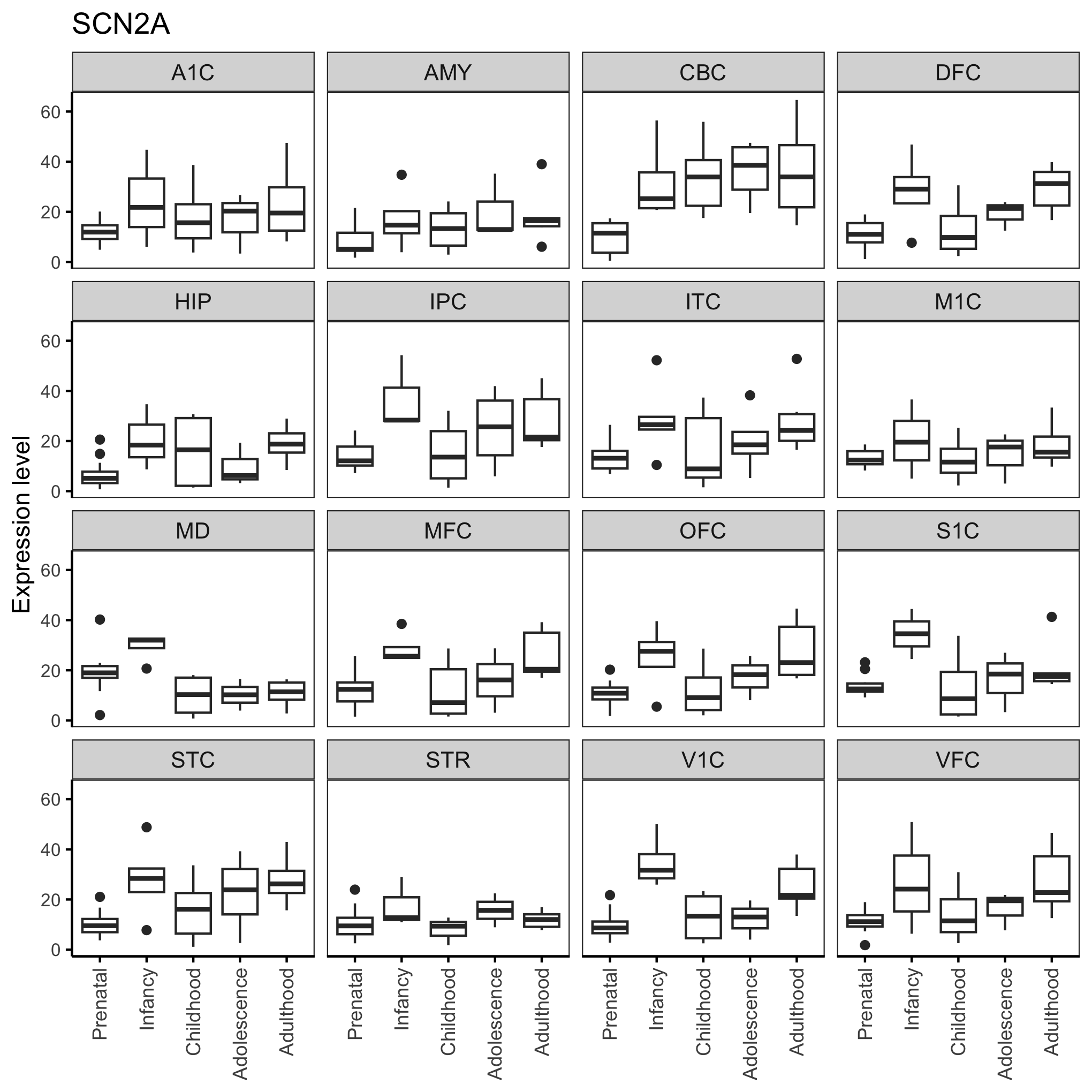

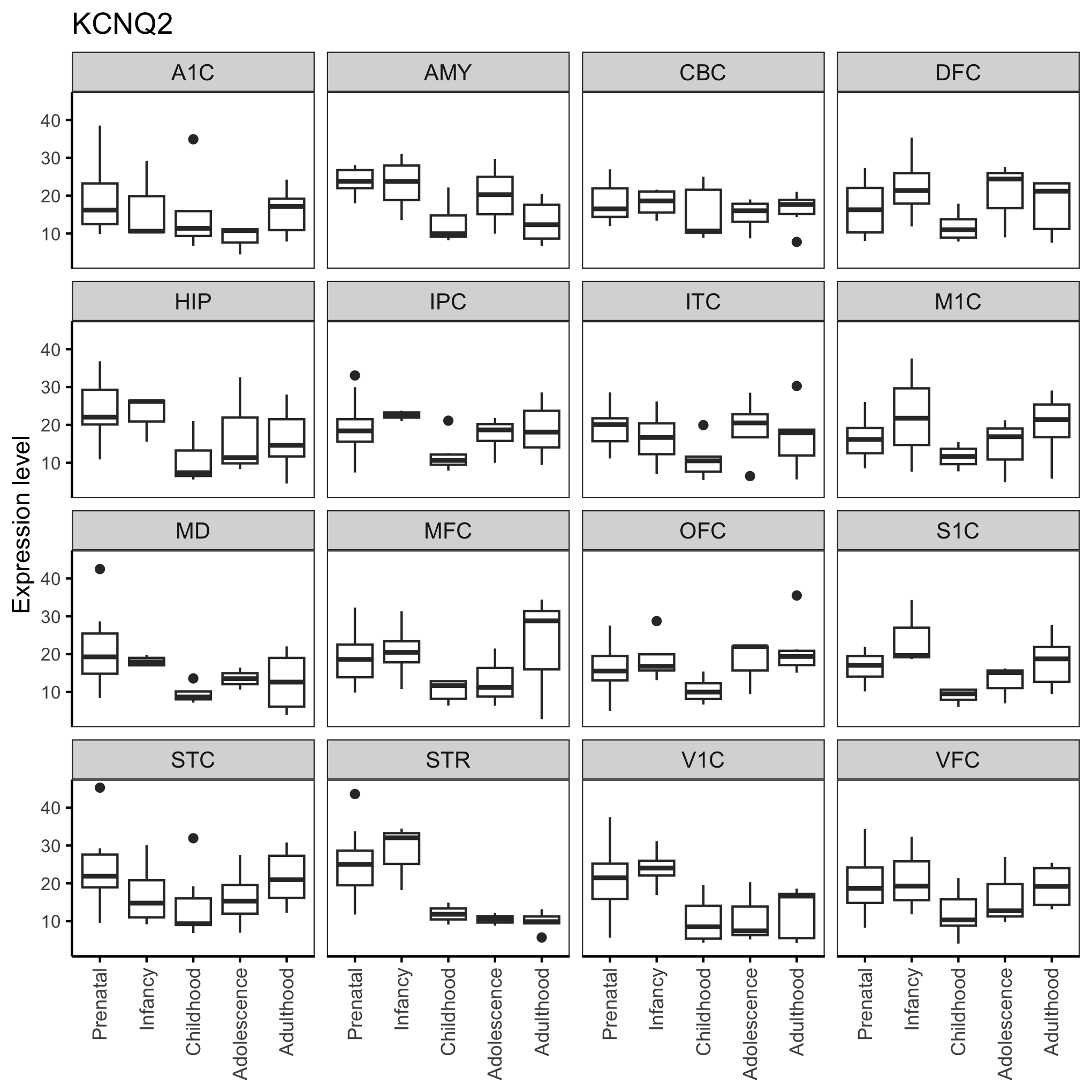

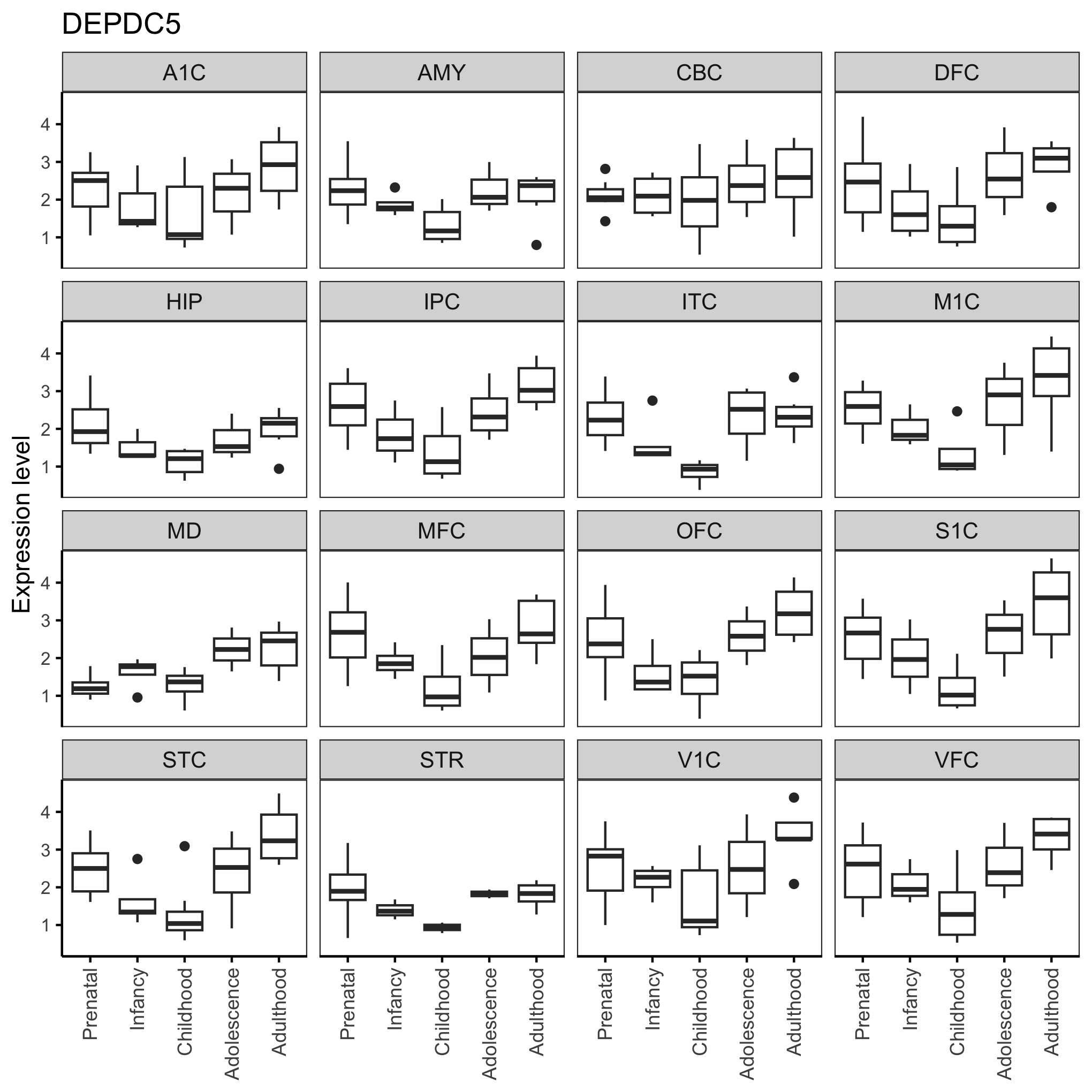

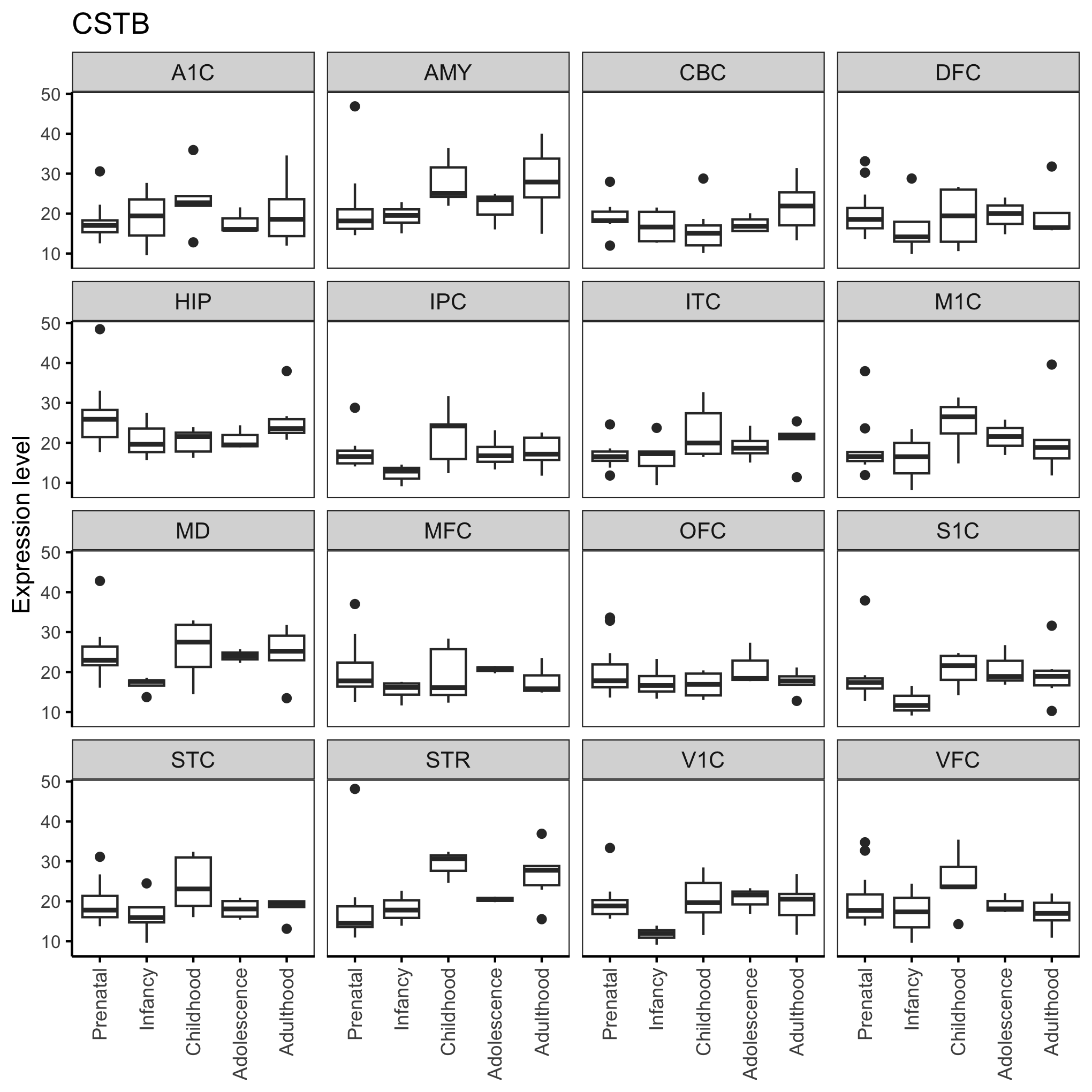

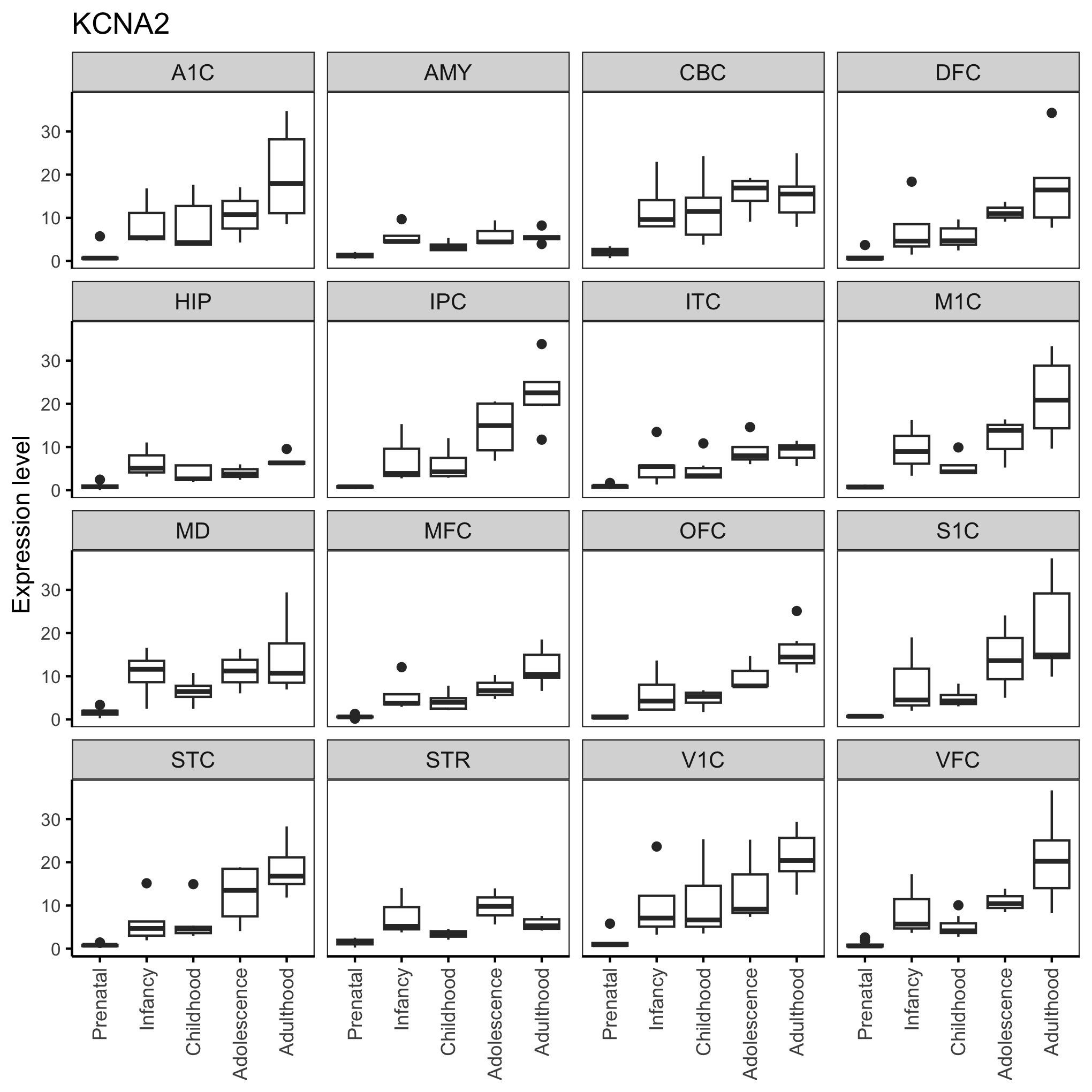

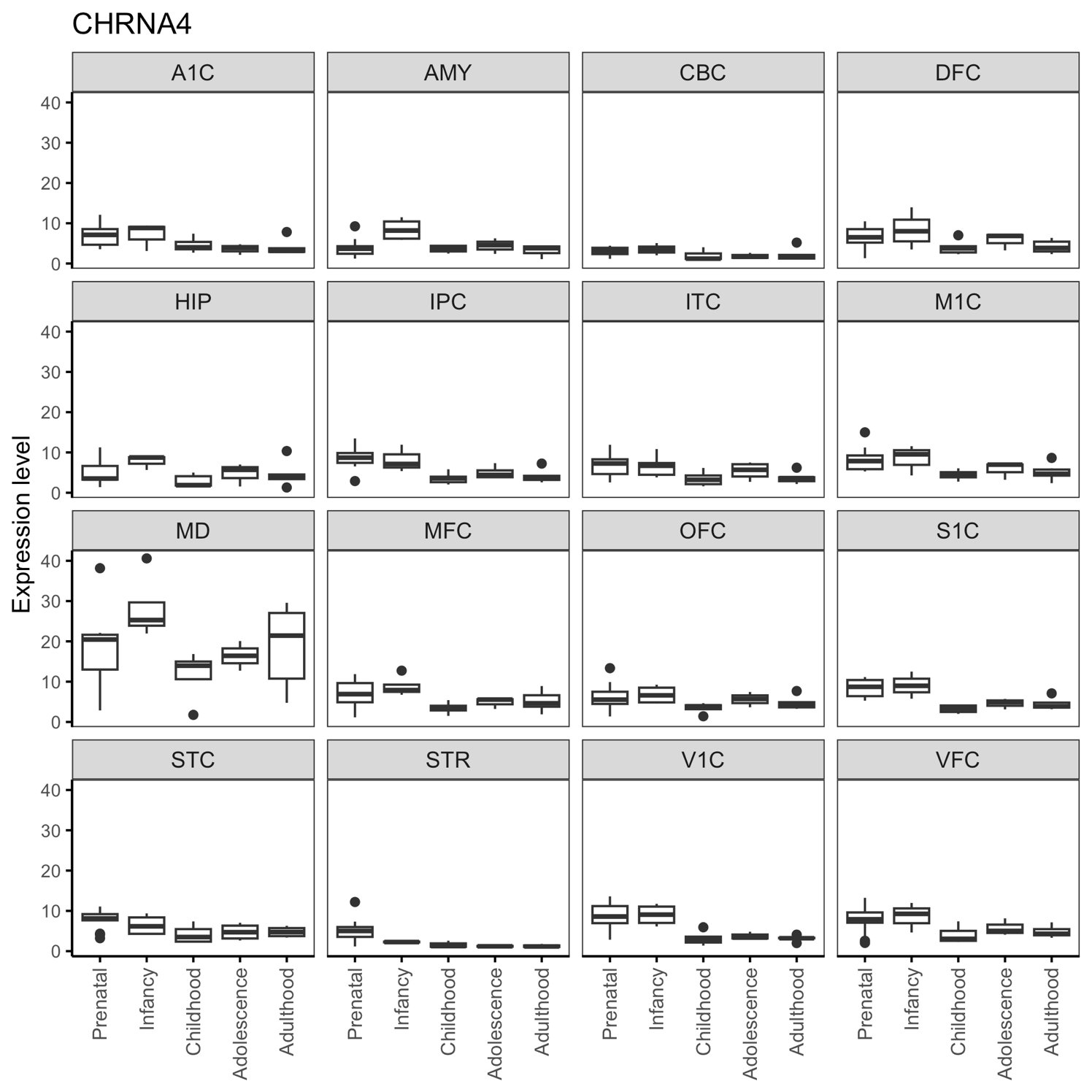

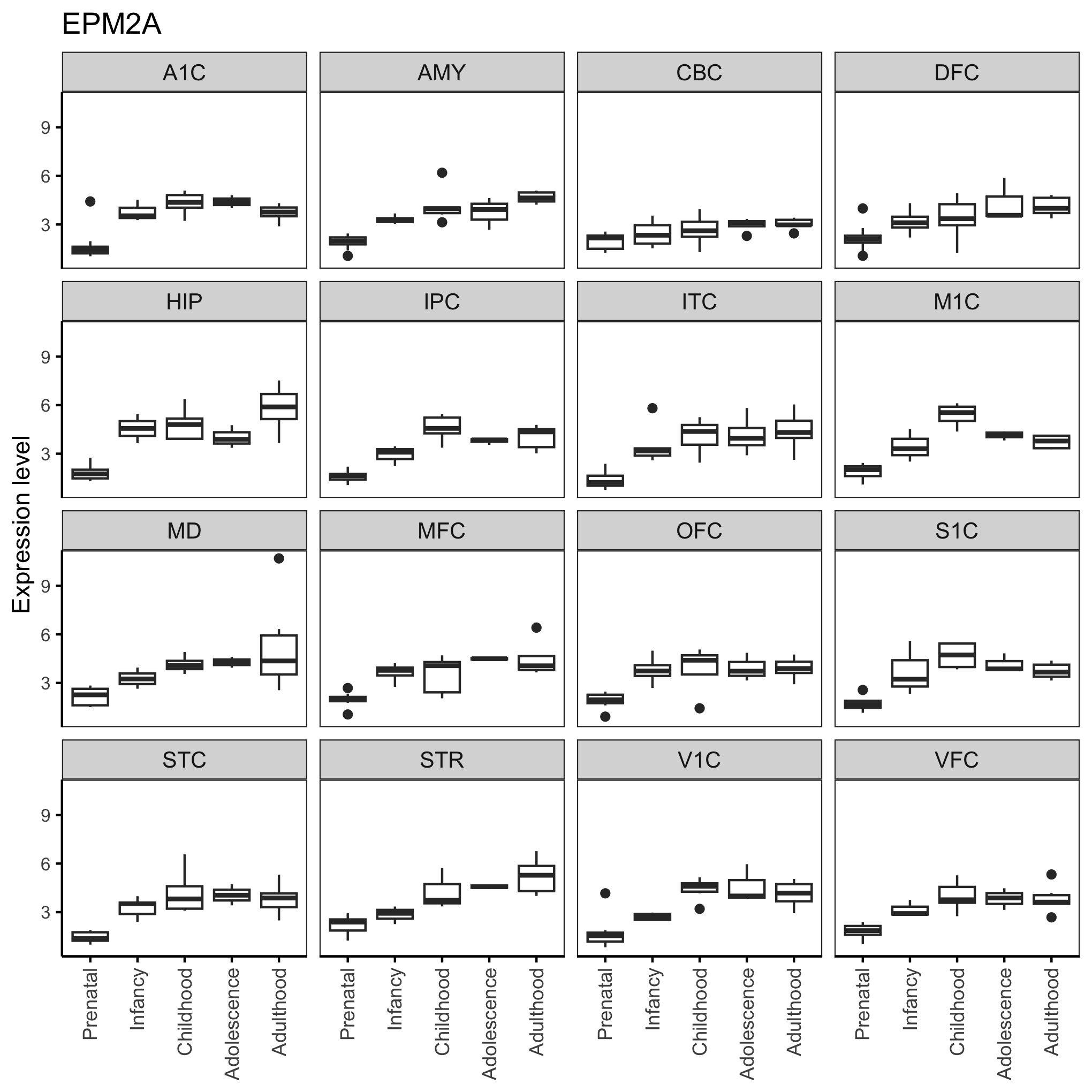

**Supplemental Figure 2. Expression of top 10 studied genes in different brain regions across development.**
