## Supplemental Fig 3 for "Integrative Analysis of Epilepsy-Associated Genes Reveals Expression-Phenotype Correlations"

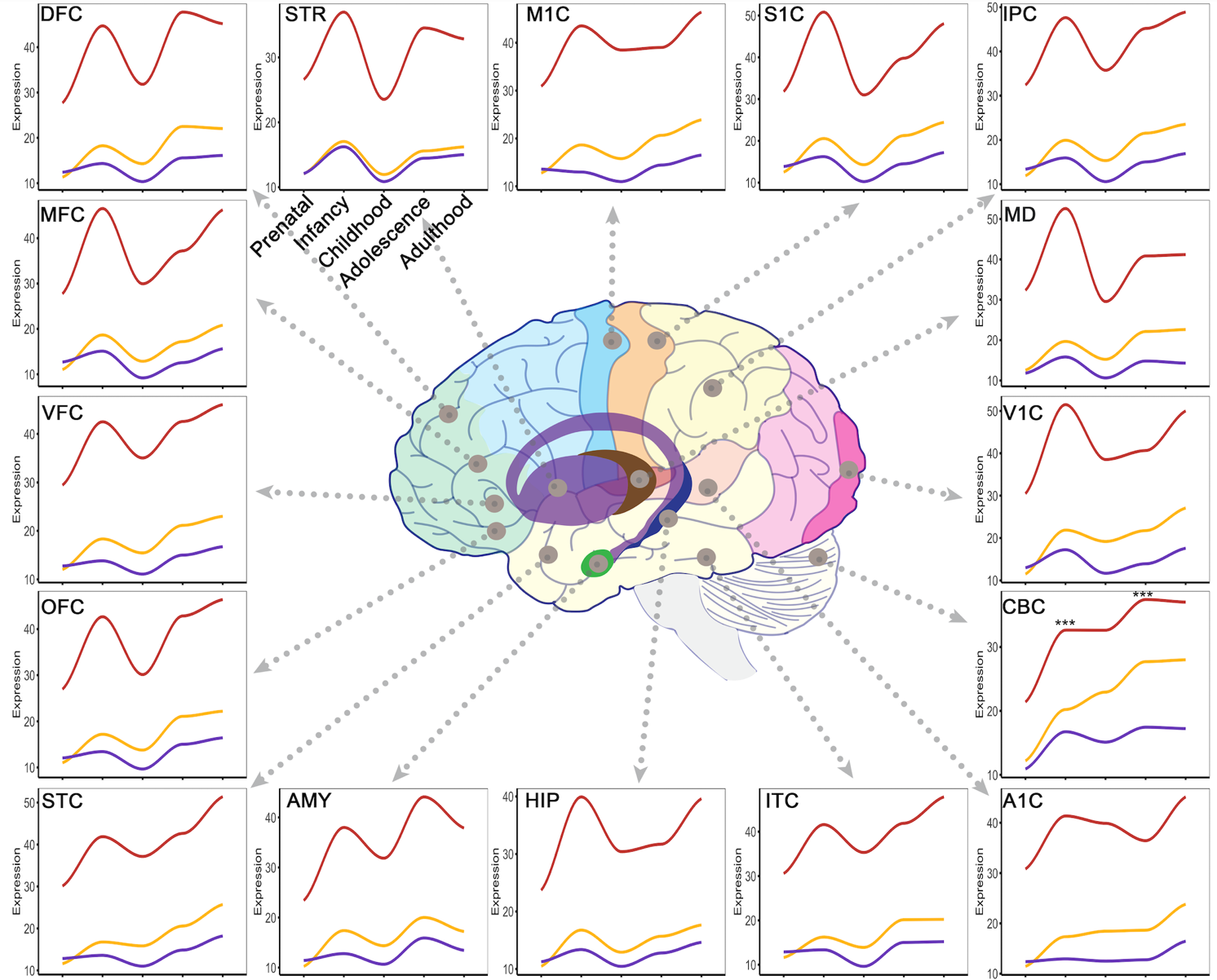

**Supplemental Figure 3. Averaged expression of epilepsy-associated genes within each group across development.** Red line: DEEG; orange: CEG; purple: SRG. Bulk RNA-seq data are from Allen BrainSpan. DFC: dorsolateral prefrontal cortex; MFC: anterior cingulate cortex; VFC: ventrolateral prefrontal cortex; OFC: orbital frontal cortex; STC: posterior superior temporal cortex; AMY: amygdala; HIP: hippocampus; ITC: inferolateral temporal cortex; A1C: primary auditory cortex; CBC: cerebellar cortex; V1C: primary visual cortex; MD: Mediodorsal nucleus of thalamus; IPC: posteroventral parietal cortex; S1C: primary somatosensory cortex; M1C: primary motor cortex; STR: striatum. See Supplemental Fig. 4 for significance levels.
