## Supplemental Fig 4 for "Integrative Analysis of Epilepsy-Associated Genes Reveals Expression-Phenotype Correlations"

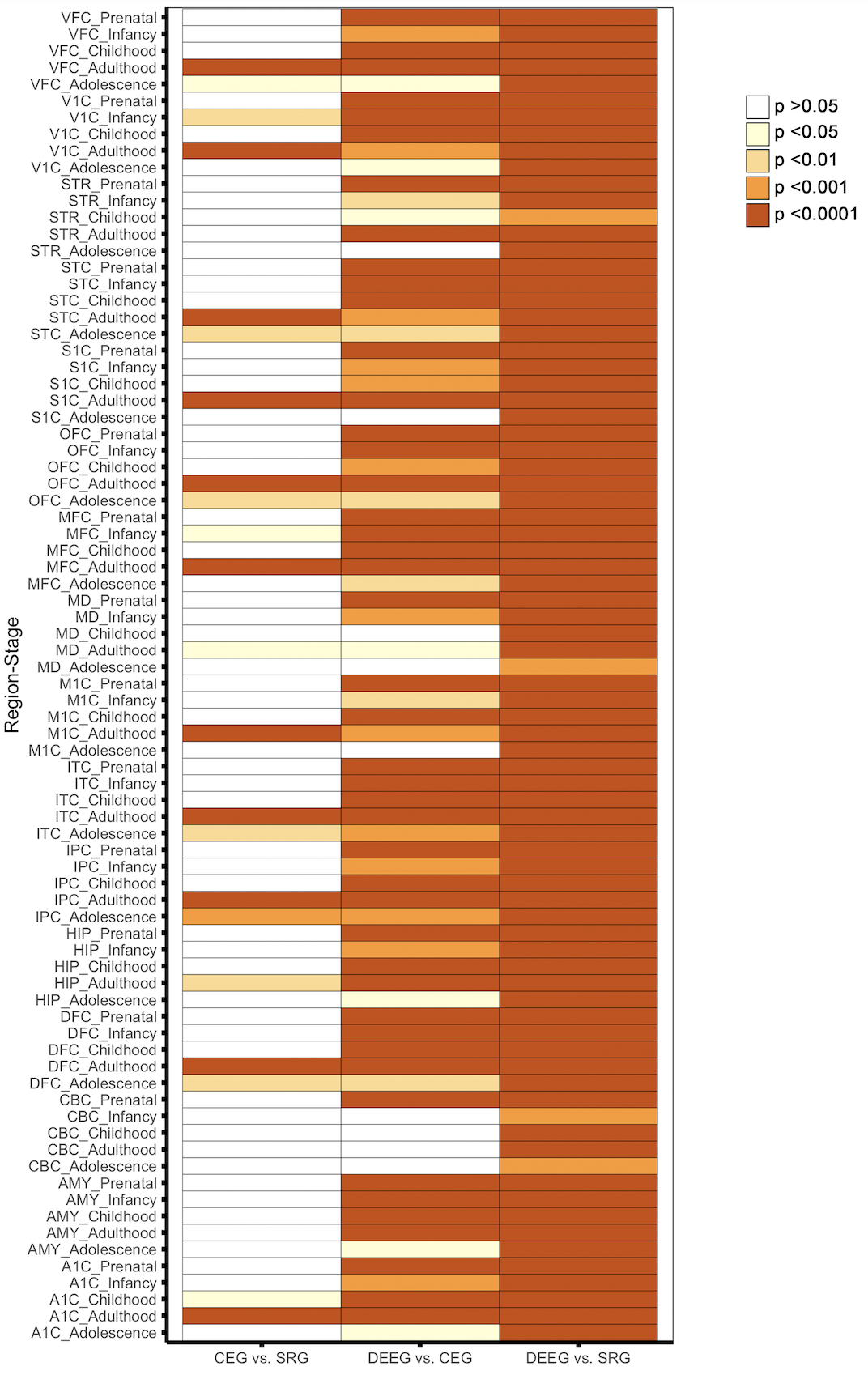

**Supplemental Figure 4. Heatmap of p-values from Wilcoxon signed rank sum test with Benjamini-Hochberg post hoc among the three groups of epilepsy-genes within each brain region for each developmental stage.** Bonferroni correction was applied for multiple tests across 16 different brain regions and 5 developmental periods. Significance levels are color coded.
