## Supplemental Fig 5 for "Integrative Analysis of Epilepsy-Associated Genes Reveals Expression-Phenotype Correlations"

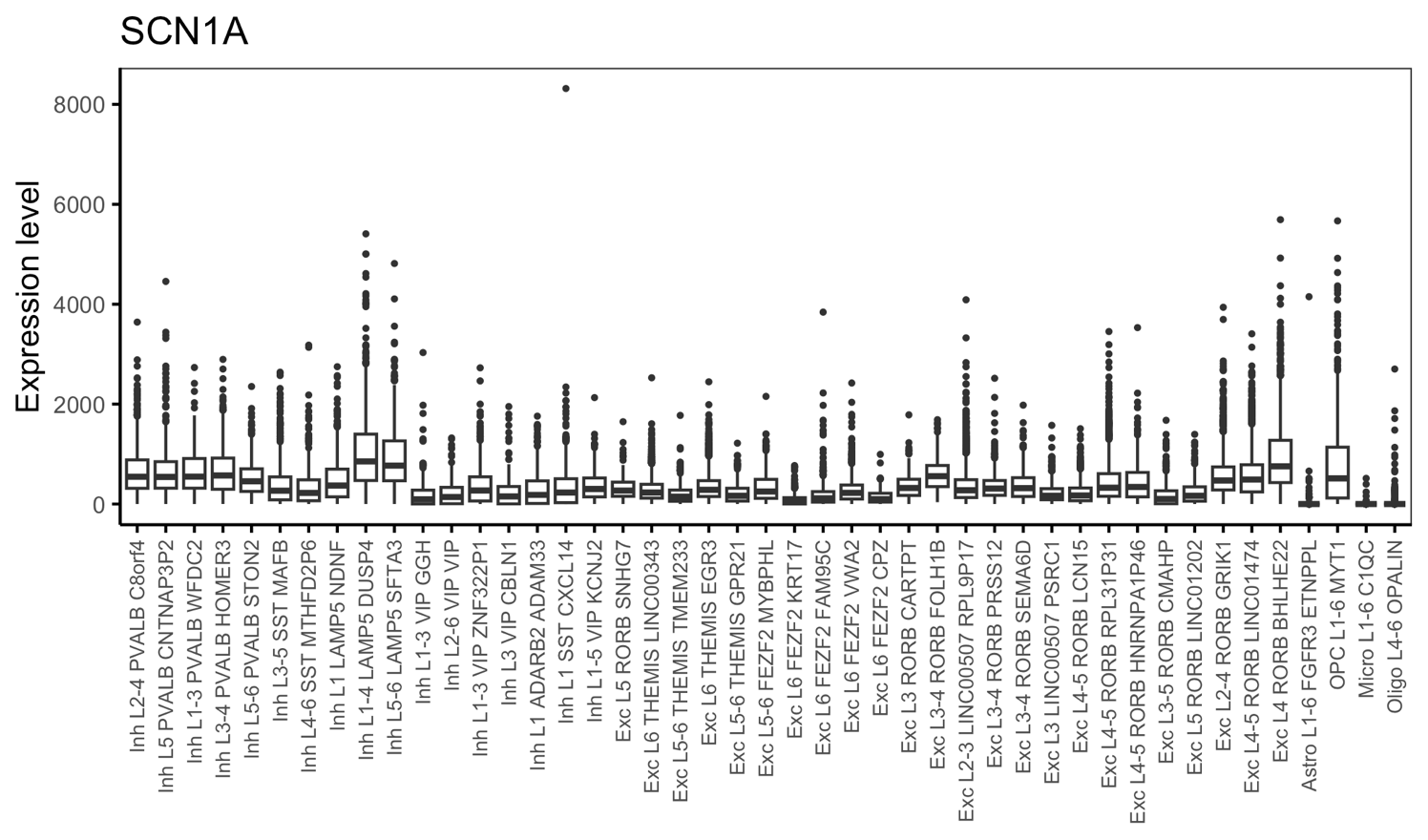

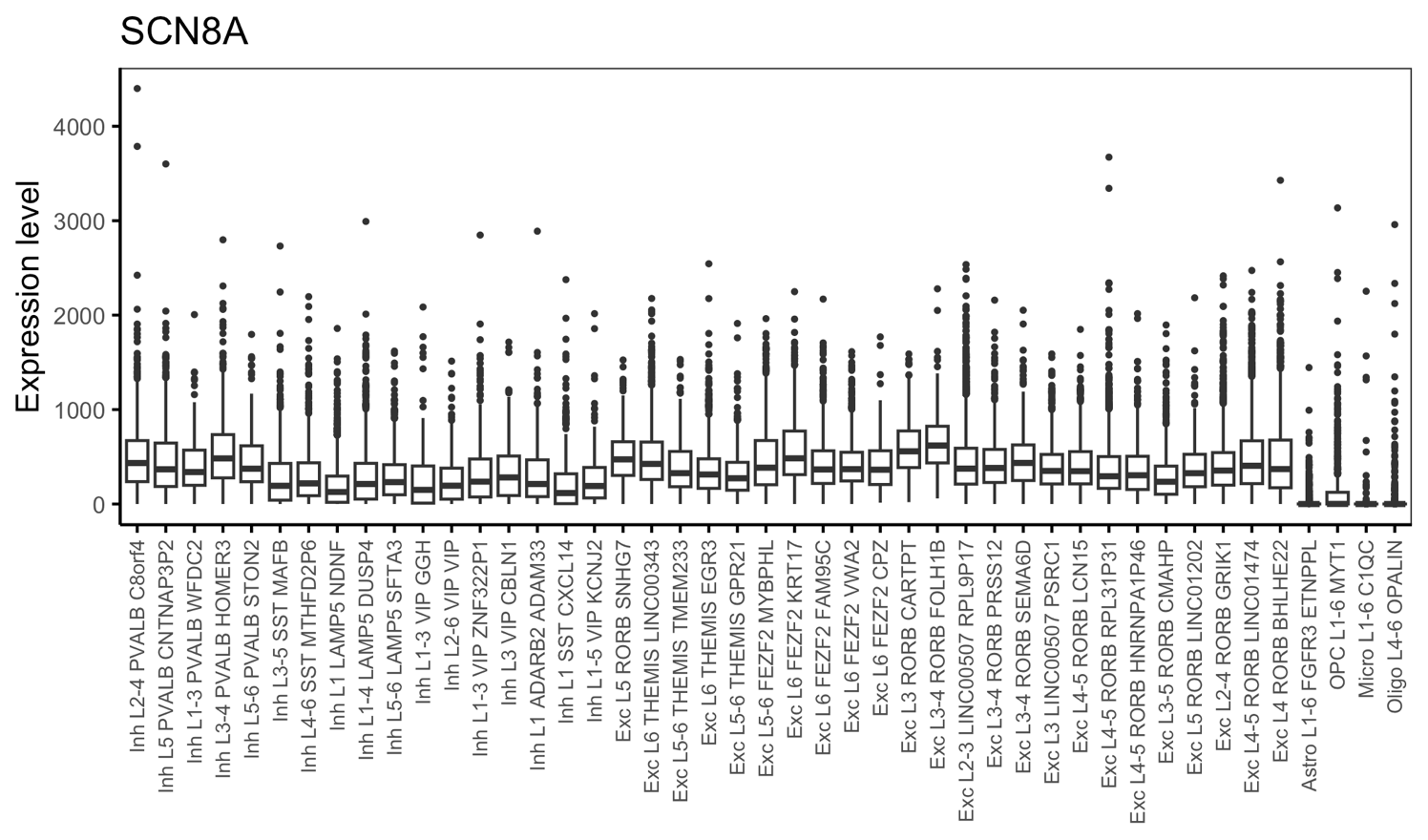

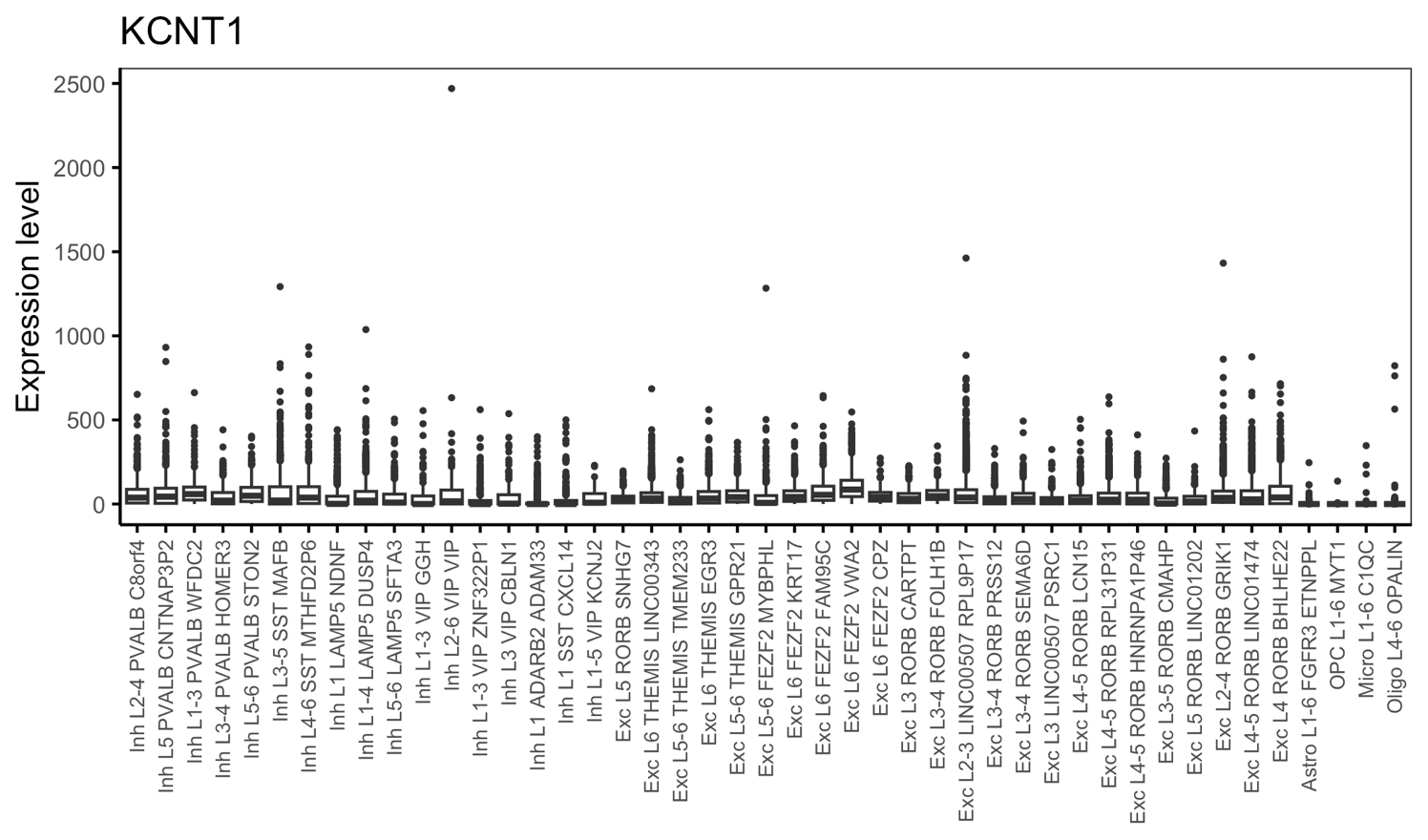

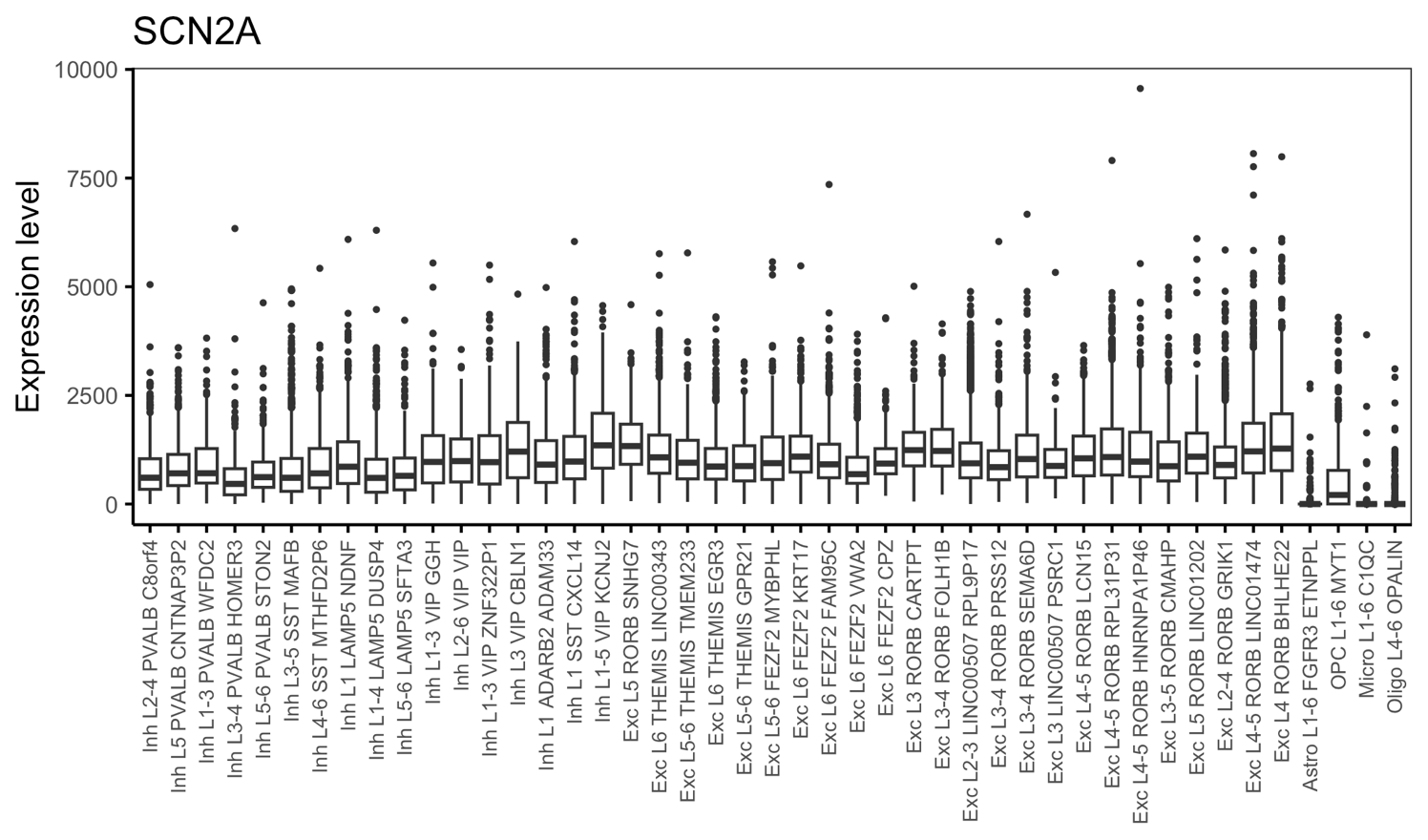

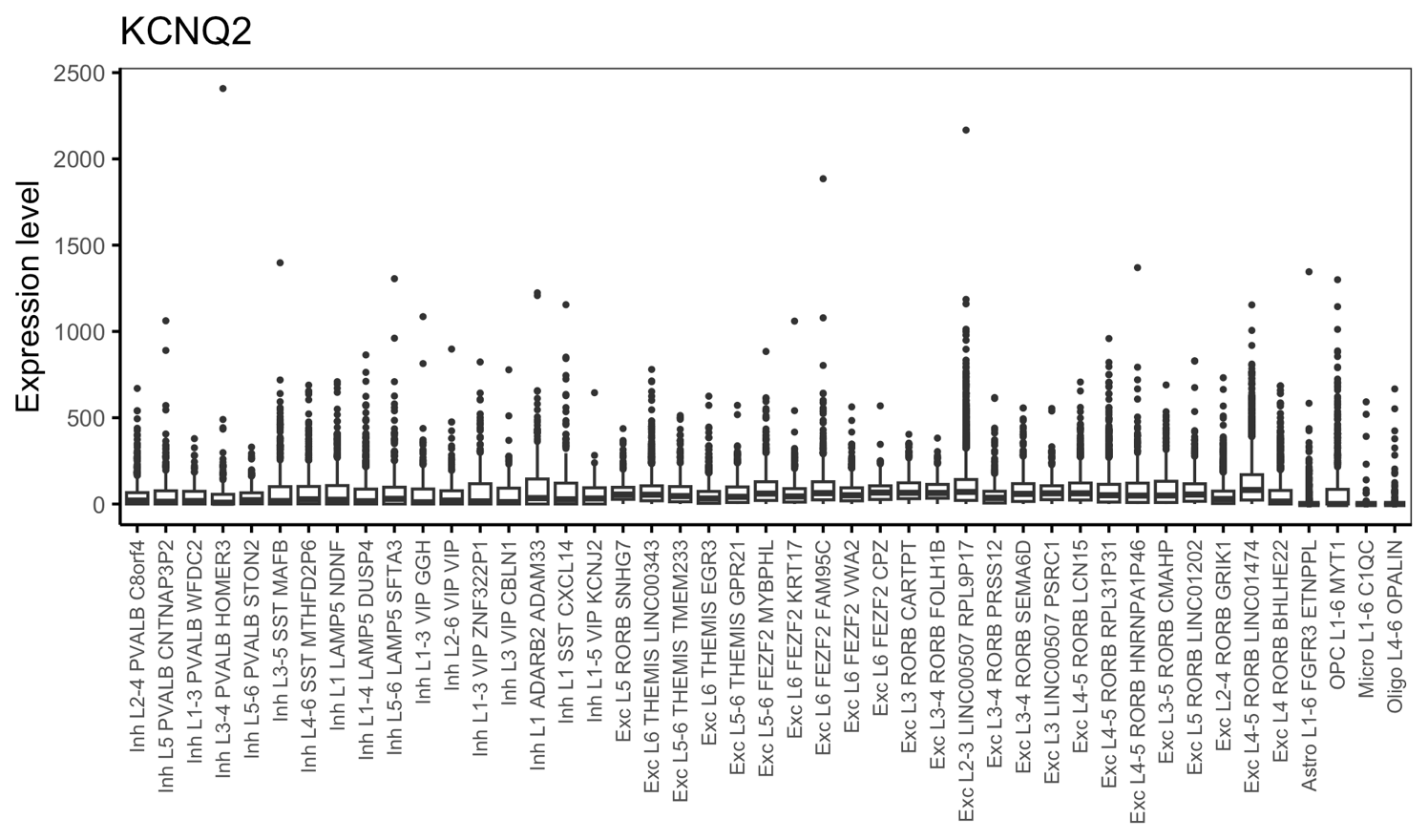

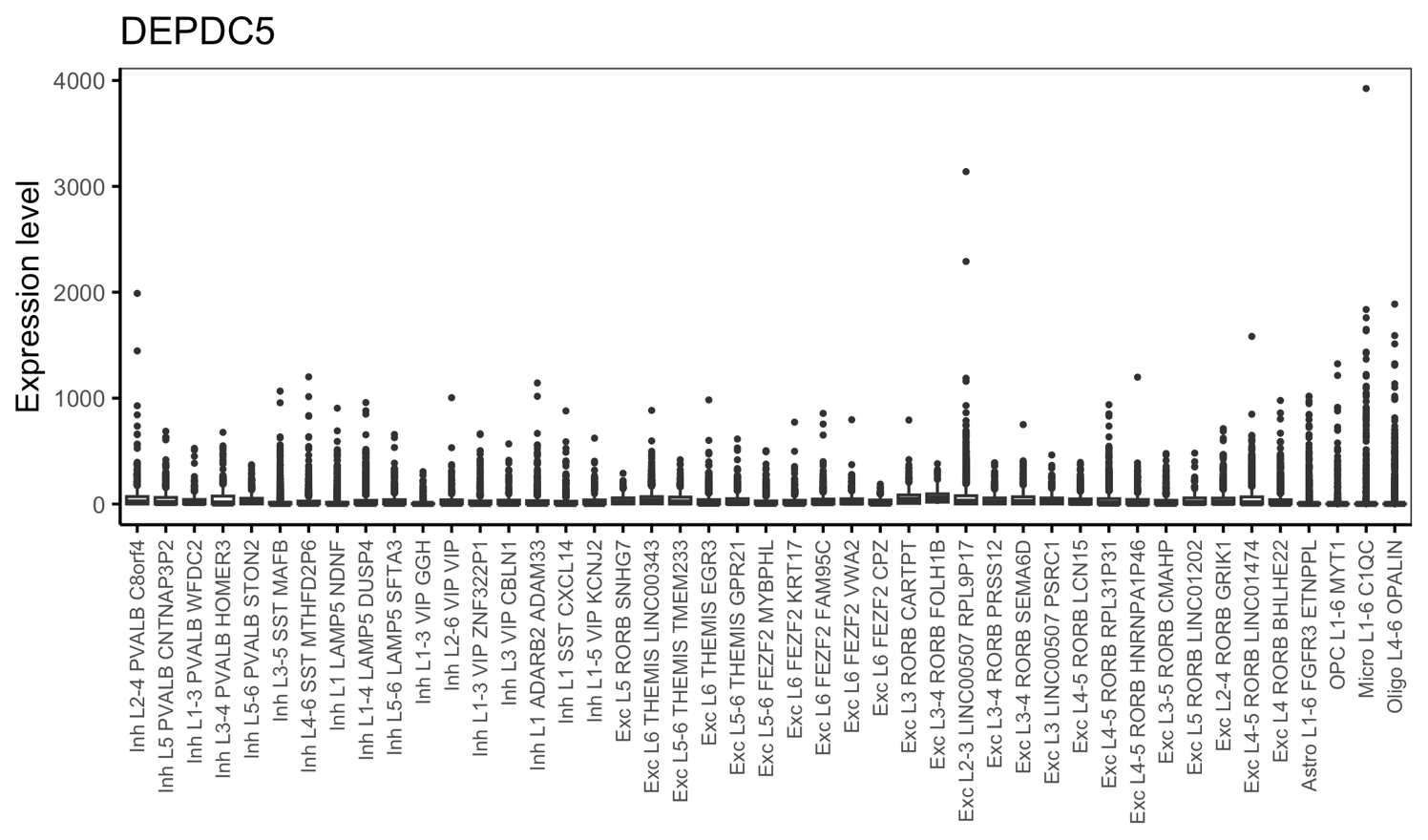

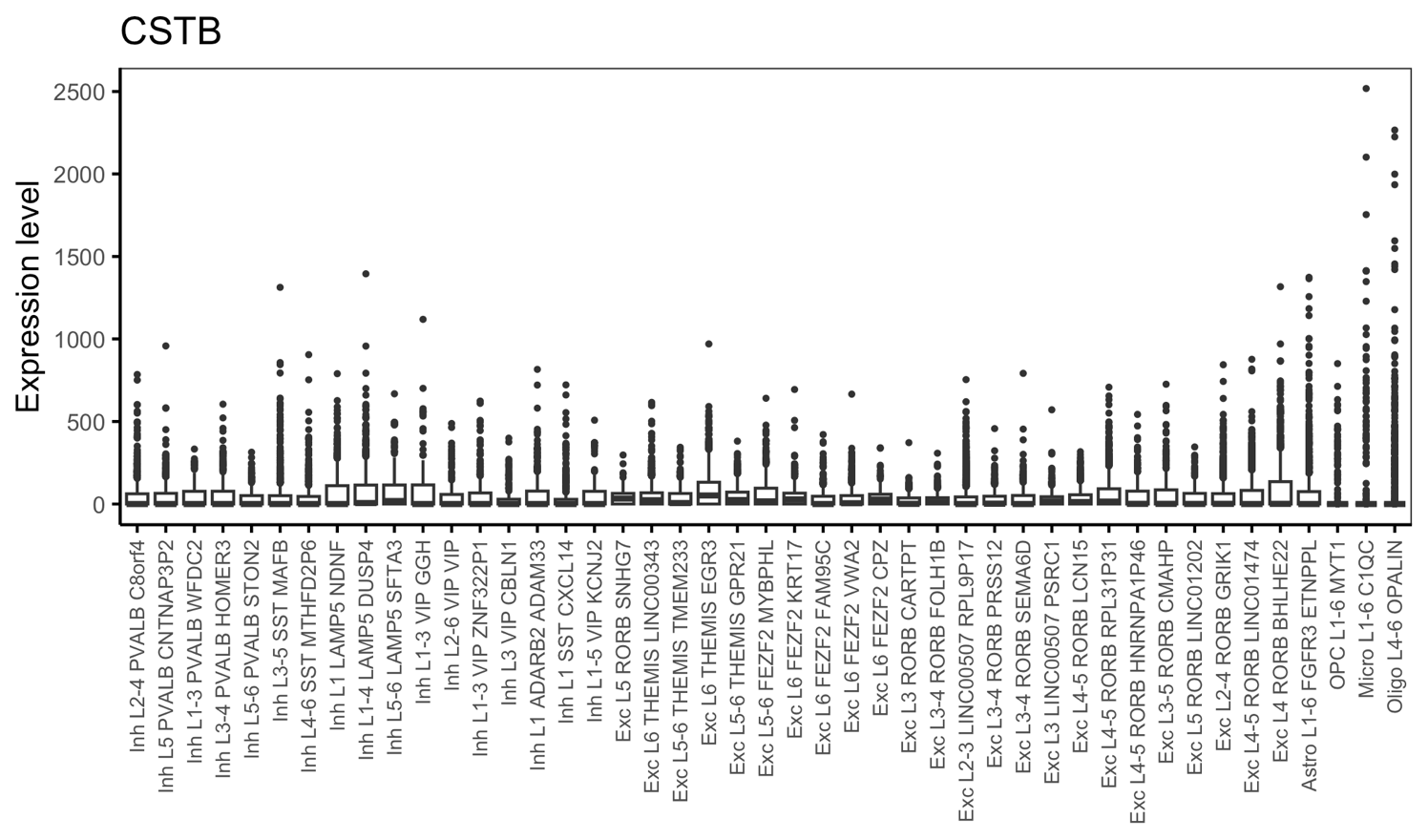

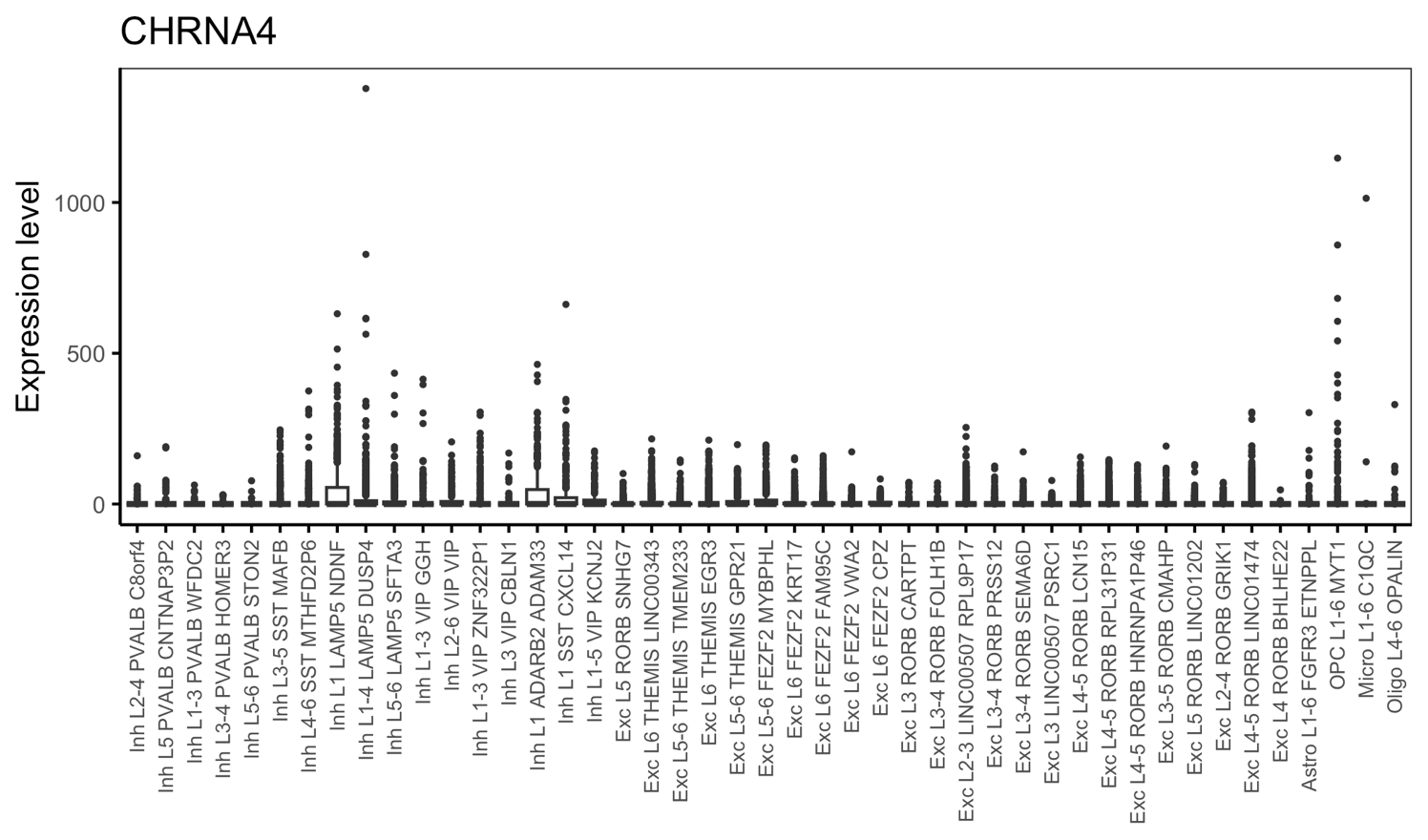

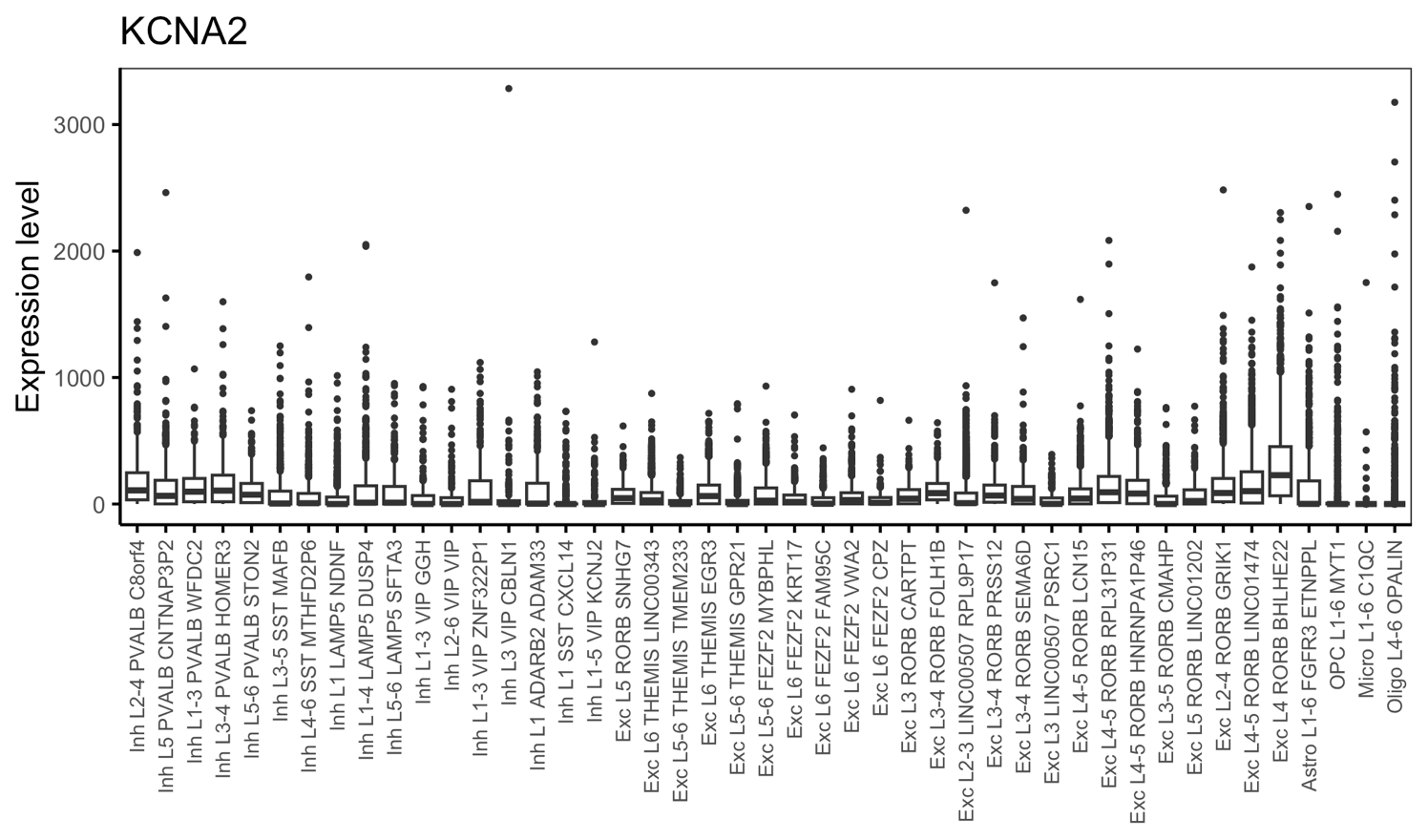

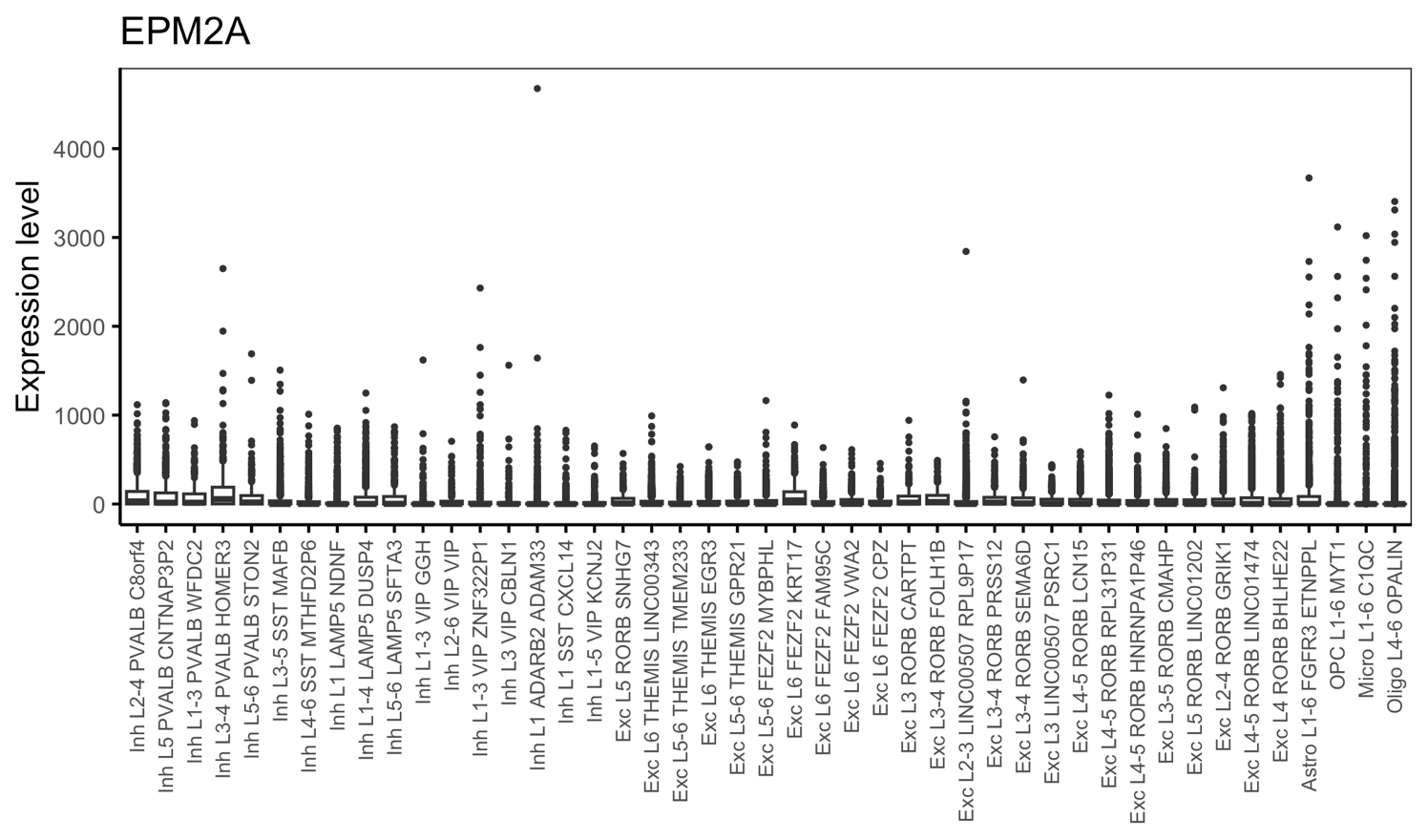

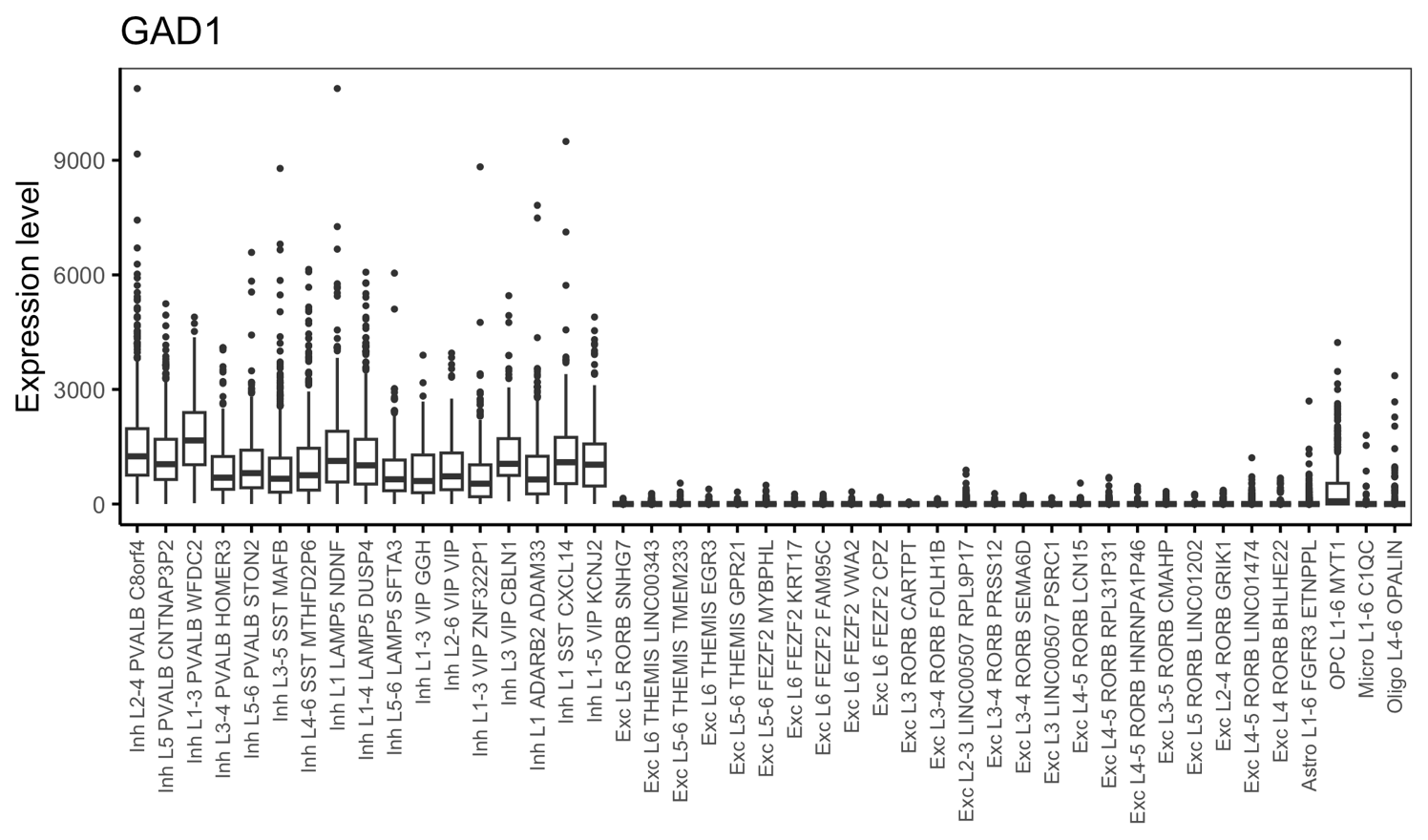

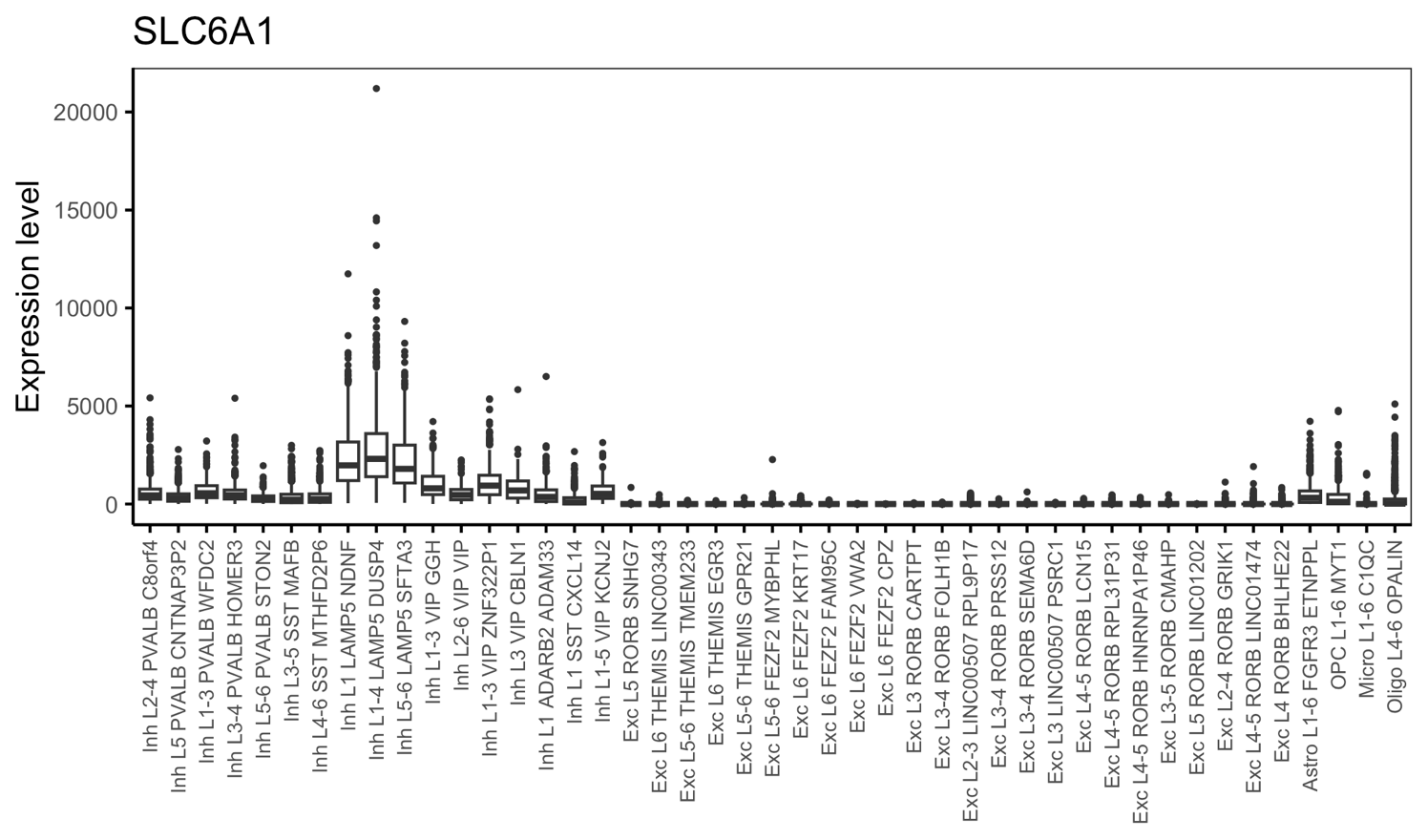

**Supplemental Figure 5. Expression of top 10 studied genes and 2 cell type marker genes in different brain cell types.**
